# Accurate strain-level microbiome composition analysis from short reads

**DOI:** 10.1101/2022.01.26.477962

**Authors:** Herui Liao, Yongxin Ji, Yanni Sun

**Author notes:** these authors contributed equally to this work (Herui Liao, Yongxin Ji).

## Abstract

Because bacterial strains can exhibit different biological properties, strain-level composition analysis plays a vital role in understanding the functions and dynamics of microbial communities. Metagenomic sequencing has become the major means for probing the microbial composition in host-associated or environmental samples. Despite a plethora of composition analysis tools, they are not optimized to address the challenges in strain-level analysis: a reference database with highly similar reference strain genomes and the presence of multiple strains under one species in a sample. In this work, we present a new strain-level composition analysis tool named StrainScan that employs a novel tree-based *k*-mer indexing structure to strike a balance between the strain identification accuracy and the computational complexity. We rigorously tested StrainScan on many simulated and real sequencing data and benchmarked StrainScan with popular strain-level analysis tools including Krakenuniq, StrainSeeker, Pathoscope2, Sigma, StrainGE, and Strainest. The results show that StrainScan has higher accuracy and resolution than the the state-of-the-art tools on strain-level composition analysis. It improves the F1-score by 20% in identifying multiple strains with at least 99.89% average nucleotide identity. StrainScan takes short reads and a set of reference strains as input and its source codes are freely available at https://github.com/liaoherui/strainScan.

## Introduction

There is accumulating evidence showing that strains within a species can have different metabolic and functional versatility due to the genomic variations^1–3^. Different strains of the same bacterium typically share 80-90% of their genes. Unique genes or SNPs to a strain may lead to new enzymatic functions, antibiotic resistance, virulence, different infecting viruses, etc. For example, there are at least thousands of strains identified for *E. coli*, with some of them containing virulence factors while others being commensal. The 2011 E. coli outbreak in Germany is caused by a strain O104:H4 that acquired a Shiga toxin-encoding prophage and other virulence factors^4^. As different strains can have different biological properties, pinpointing the strains is not only important for high-resolution analysis of the composition, but also important for downstream applications such as precise diagnosis and treatment.

Metagenomic sequencing data, which contain sequenced genetic materials from a host-associated or environmental sample, become a major source to study strain-level compositions in the underlying samples. Strain-level analyses of bacteria have been applied to a variety of microbiomes in order to achieve a more precise understanding of the relationship between the genotype and phenotype. For example, it showed that many prevalent bacterial species have stain-level composition associated with a geographic location in 198 marine metagenomes^5^. A closely related study showed that dominant E. *coli* strains change over time in the gut microbiome of a Crohn’s disease patient^6^. *P. copri*, another very common bacterium in the human gut, has been proven to have a tight link between its strains and the host’s geographical location and dietary habits^7, 8^. Some strains of the potential probiotic *A. muciniphila* are found to have anti-inflammatory properties, which could have beneficial effects on obesity and diabetes^9^. In addition, there are differences in the distribution of strains in different parts of the human body. For example, a past study^10^ has found that strains of *C. acnes* and *S. epidermidis* collected from different sites of the body are heterogeneous and multiphyletic.

Despite the importance of strain-level analysis, it has been difficult to conduct the taxonomic analysis below the species level. One challenge comes from the fact that multiple strains can exist simultaneously in one sample. For example, there are reports showing that multiple strains of *C. acnes*, an important component in the human skin microbiome, often form a complex mixture^11^. Some of these strains may share high sequence similarities (>99% sequence similarity). Commonly used metagenomic binning and assembly tools are not designed to distinguish different strains. Although there are strain-analysis tools, they may either require multiple samples from the same population^12^, only output the dominant strain^13–16^, or pose a restriction on the similarity of between the strains^17^. The second challenge that immediately follows is the resolution of strain-level identification. The resolution here is reflected by the size of the reference database, with a larger number of reference strains indicating a higher resolution^18^. A higher resolution is usually preferred because it gives us a more complete picture of the true strain composition of the sample. Tools including StrainGE19 and Strainest^20^, are designed to untangle strain mixtures, but are limited to reporting a representative strain in a sampled strain database, and are unable to distinguish distinct strains that are close to the same representative strain. Two *k*-mer-based tools, Krakenuniq21 and StrainSeeker^22^, also have a very low resolution in strain-level identification when strains in the database share high similarities. The third challenge is the identification of low-abundance strains. For example, the de novo strain construction tools^23, 24^, which aim to reconstruct strains by using assembly-based strategies, usually require a high coverage of strains to achieve an accurate strain reconstruction. Besides, many strain-analysis tools^25–27^ also require strain coverage greater than 10X to return accurate identification. Thus, it remains a challenge to identify strains with low coverage for these tools. The last challenge is the strain identification time. According to the recently published studies^18, 28^, most alignment-based strain-level identification tools including Sigma^29^, Pathoscope230 can be computationally expensive when the database is large. While the large reference database can increase coverage of intra-species diversity, it also requires more computational resources.

Thus, there is a pressing need to provide more sensitive, accurate, and efficient strain-level analysis for metagenomic data. In this work, we introduce StrainScan, an open-source tool that can accurately detect known strains from sequencing data, including metagenomic data or whole-genome sequencing data. In order to strike a balance between the resolution and computational complexity, we developed a novel hierarchical *k*-mer indexing structure for a large number of strains, which usually demonstrate heterogeneous similarity distribution. In the first step, we cluster highly similar strains into clusters. Then we design a novel Cluster Search Tree (CST), which is a tree-based indexing structure for cluster search. CST is a binary tree and each node contains sufficient *k*-mers representing strains in its rooted sub-tree. By carefully balancing the number of *k*-mers in each node, we optimize the CST to prevent false positive strain identification for low abundance strains. In the second step, we use strain-specific *k*-mers and *k*-mers that represent SNVs and structural variations to determine which strains are likely to present. The final output of StrainScan includes the identified strains and their abundance.

By benchmarking StrainScan with other available tools on multiple simulated and real sequencing datasets, we demonstrate that StrainScan can output strain-level composition with higher accuracy than the state-of-the-art tool. In particular, when compared to high-accuracy tools such as Strainest, StrainScan improved the F1-score by more than 20% in identifying multiple strains with at least 99.89% average nucleotide identity and increased the speed by at least one magnitude. StrainScan is a targeted strain composition analysis tool, requiring the users to provide reference genomes for bacteria of interest. It can be applied to any bacteria because it supports the construction of the indexing structure for any given set of reference genomes.

## Results

Because StrainScan focuses on identifying known strains, we test the performance of StrainScan on different bacteria that tend to pose computational challenges for strain-level analysis. All chosen bacteria have at least 100 sequenced strains and they are listed in Figure 1A. Some of them have a large number of known strains such as *E. coli* and *S. epidermidis*. Some have strains with extremely high sequence similarity, such as *M. tuberculosis*. In addition, we choose bacteria that usually inhabit different ecosystems such as the human gut and human skin. We carried out multiple experiments to evaluate StrainScan. First, we tested the ability of StrainScan in identifying one strain and multiple co-existing strains in simulated data. We generated different datasets by configuring the parameters such as strain similarity and strain sequencing depth, which helps us compare the performance of different tools in difficult scenarios. Second, we tested StrainScan in three mock community datasets, which allow us to evaluate different tools in real sequencing data with known strain composition. Third, we tested StrainScan in 57 real sequencing datasets. Because there is usually no ground truth for the stain composition in the real sequencing data, we choose the datasets that had been analyzed by other strain identification tools^16, 20, 28, 30^. By comparing the analysis results, we are able to draw some conclusions about different tools’ performance. In these experiments, we used the F1 score, precision, recall, and Jensen–Shannon divergence as the evaluation metrics, which were defined in Methods. We benchmarked StrainScan against popular reference-based strain-level analysis tools including Krakenuniq^21^, StrainSeeker^22^, Pathoscope2^30^, Sigma^29^, StrainGE^19^ and Strainest^20^.

**Figure 1.**
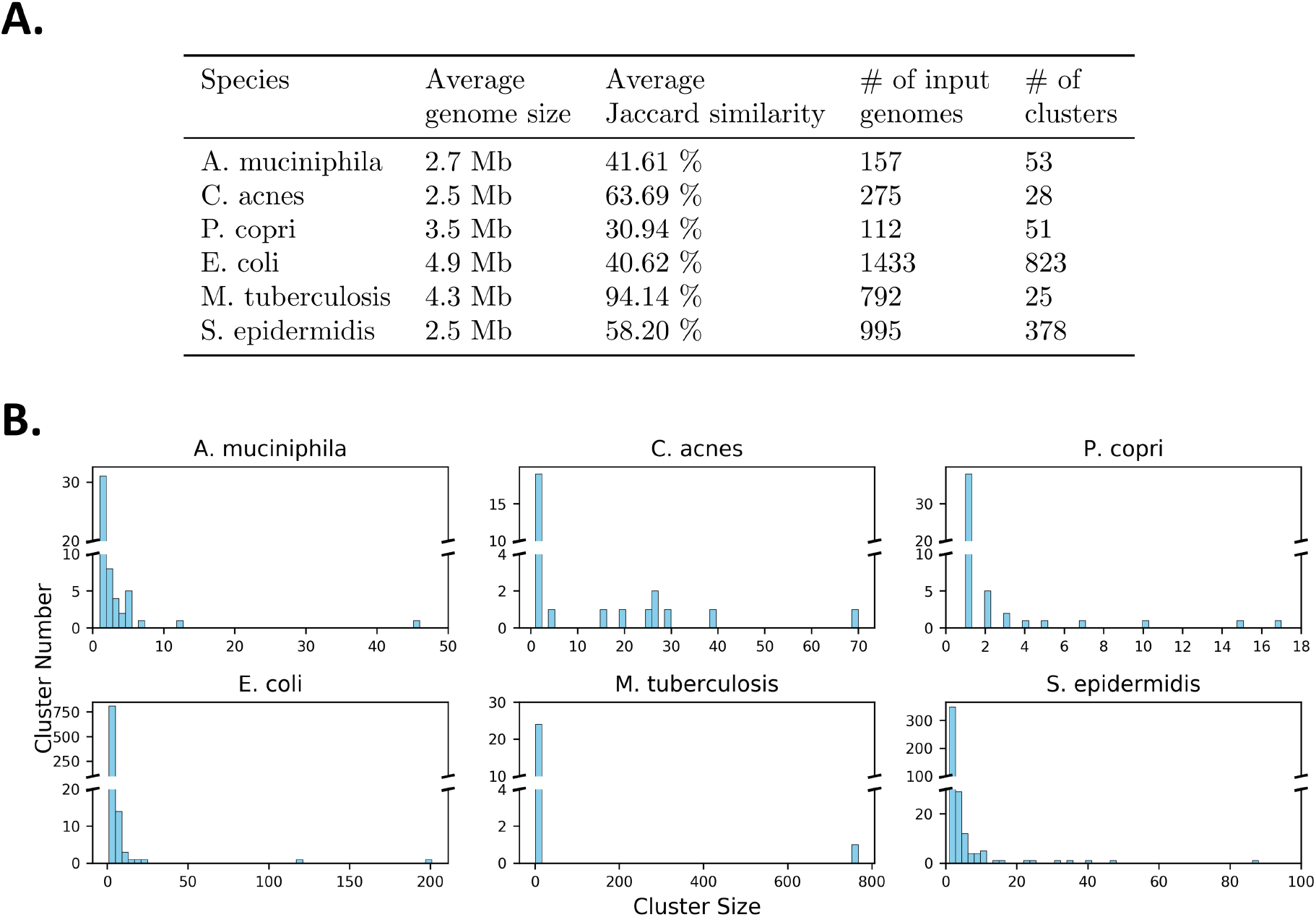
A. The summary statistics of the reference genomes for 6 representative bacterial species. “Average Jaccard similarity” is obtained by calculating the average of *k*-mer Jaccard similarity of all strains using Dashing^31^. B. The histogram of cluster sizes for clusters in the StrainScan’s reference databases.

The output of all the tools is a list of strains ranked by confidence. Thus, for experiments on the simulated data that contains *x* strains, we will keep only the top *x* outputs of a tool when calculating related metrics. The false positive (FP) cases can be contributed by different types of outputs because of the usage or output formats of different tools. We carefully define three FP cases: 1) The tool outputs a strain that is not in the ground truth (i.e. not present in the sample, see Case2 in Supplementary Figure S6). 2) The tool outputs multiple strains with the same score with one of them being correct, such as the case of StrainSeeker (see Case3 in Supplementary Figure S6). In this case, only one strain is regarded as TP and others are FPs. 3) Tools such as StrainGE and Strainest group reference genomes into clusters and select one representative strain to represent all strains in the same cluster. As a result, they do not intend to further distinguish strains in the same cluster. We examine whether the representative strain is in the ground truth. If not, it is regarded as FP. (see Case5 in Supplementary Figure S6). However, if the cluster of identified representative strain doesn’t contain the ground truth, then it will be treated as case 1). Below we present the experimental results.

### Reference database construction

For all the species tested in this work, we created the reference strain genome database as comprehensively as possible. Thus, we downloaded all complete and draft genomes from the NCBI RefSeq database for the tested bacteria. But there are 25,349 *E. coli* genomes, requiring >1TB memory. Due to the constrain of our hardware resources, we only used the complete E. *coli* genomes from RefSeq. Similar to E. *coli*, our hardware resources prevent us from using all draft and complete genomes for *M. tuberculosis*. In addition, some available genomes for M. *tuberculosis* only differ by < 10 positions^28^. These near-identical strains will be clustered in our pre-processing step. Thus we computed pairwise Jaccard similarities of all *M. tuberculosis* strains using Dashing^31^ and performed complete-linkage clustering using a *k*-mer Jaccard similarity threshold of 99%. Then, we selected the strain with the highest average similarity to all other genomes in that cluster as the representative strain. Finally, we sampled 792 representative genomes out of 6,752 genomes.

The final numbers of the strains and their other properties were recorded in Figure 1A. We also show the number of clusters (Figure 1A) and the size distribution of the clusters (Figure 1B) used in the CST tree. The result shows the intra-species similarity of different bacteria have different distributions. Many bacteria have a large number of small clusters containing just 1 or 2 strains, indicating that their similarity is not very high. On the other hand, some bacteria have big clusters with highly similar strains. For example, *M. tuberculosis* has one cluster containing 768 strains. According to the algorithm described in Methods, StrainScan first locates the cluster using the CST tree and then identify the strain inside the cluster. Our experimental results show that the cluster search using CST can achieve 100% accuracy for all tested bacteria. Once pinpointing clusters, StrainScan will search strains in the identified clusters and predict their abundance using the elastic net model. For a fair comparison, the strains used to construct the StrainScan’s reference databases were also used to construct databases for all other tools. But StrainGE and Strainest will further cluster the input strain genomes and only keep one representative strain selected from each cluster in their final databases. As a result, there are significantly few strains left in their database (Supplementary Figure S2 and S3), which leads to a decrease in the resolution.

### Detecting a reference strain from simulated reads

The purpose of this experiment is to test StrainScan and other tools on identifying the correct strain in a sample. The challenge is to distinguish the true strain from other highly similar peers. As Sigma and Pathoscope2 are computationally expensive, we were not able to construct their databases for *E. coli, S. epidermidis*, and *M. tuberculosis*.

For each bacterium, we randomly picked a reference strain and used its simulated short reads as input to all tools. In order to avoid any data-related bias, we repeated the experiment 60 times, with a strain randomly picked each time. Thus, there were 360 datasets in total. For each dataset, we simulated Illumina reads using ART^32^ with the following parameters: -p -l 250 -f 10 -m 600 -s 150. StrainScan and the other six programs were used to identify strains from these simulated reads.

The F1 score, recall, and precision of each program were shown in Figure 2A. Of the 60 strains, some of them have more than 99.5% *k*-mer-based Jaccard similarity with at least one other reference strain genome. As a result, we noticed low F1 scores in several tools. StrainScan achieves perfect F1 scores on all datasets. While Pathscope2 and Sigma also achieved very high F1 scores, they were at least 10 times slower than StrainScan (Figure 2C). In addition, there are some other limitations with these two tools. As mentioned before, Sigma and Pathoscope2 were not able to construct custom databases for large-scale genomes, which made their usage limited in real scenarios. StrainGE and Strainest, only returned the representative strains from a sampled database and therefore have a lower precision than other tools. The major problem of StrainSeeker is the multiple hits with identical scores, which also leads to low precision. Intuitively, if a tool outputs multiple strains with the same score, it is difficult for users to know the actual strain present in the sample. Thus, we further investigated the number of strains falsely identified by StrainSeeker, StrainGE, and Strainest (Figure 2B). The result shows that all these three tools falsely identified more than 50 strains in some cases, which means these tools can only inform users that the correct strain is in these 50 strains but not exactly which strain it is. Furthermore, there are still a large number of SNVs between the actual strains and the representative strains in the database of StrainGE and Strainest, which may imply different biological properties of these strains (see Supplementary Figure S4). In addition, StrainSeeker and StrainGE are not able to identify strains with high *k*-mer Jaccard similarities such as those of *M. tuberculosis*. Krakenuniq achieves a higher F1 score for the datasets that many strains have unique *k*-mers. However, high similarity strains lead to low recall for Krakenuniq on some bacteria. As shown in Figure 2C, StrainScan is efficient in all tested bacteria except *M. tuberculosis*. Due to high *k*-mer-based Jaccard similarities across strains of *M. tuberculosis*, StrainScan assigned most of the strains to one big cluster with a significant number of *k*-mers (Figure 1B), and thus StrainScan took more time to identify them. Nevertheless, StrainScan still has the best performance in terms of the identification of *M. tuberculosis* strains. All the experiments were tested on an HPCC CentOS 6.8 node with 2.4Ghz 14-core Intel Xeon E5-2680v4 CPUs and 128 GB memory. We used 8 threads for all tools. In summary, StrainScan is able to achieve higher precision without sacrificing resolution, even when the true strain has peers of high sequence similarity.

**Figure 2.**
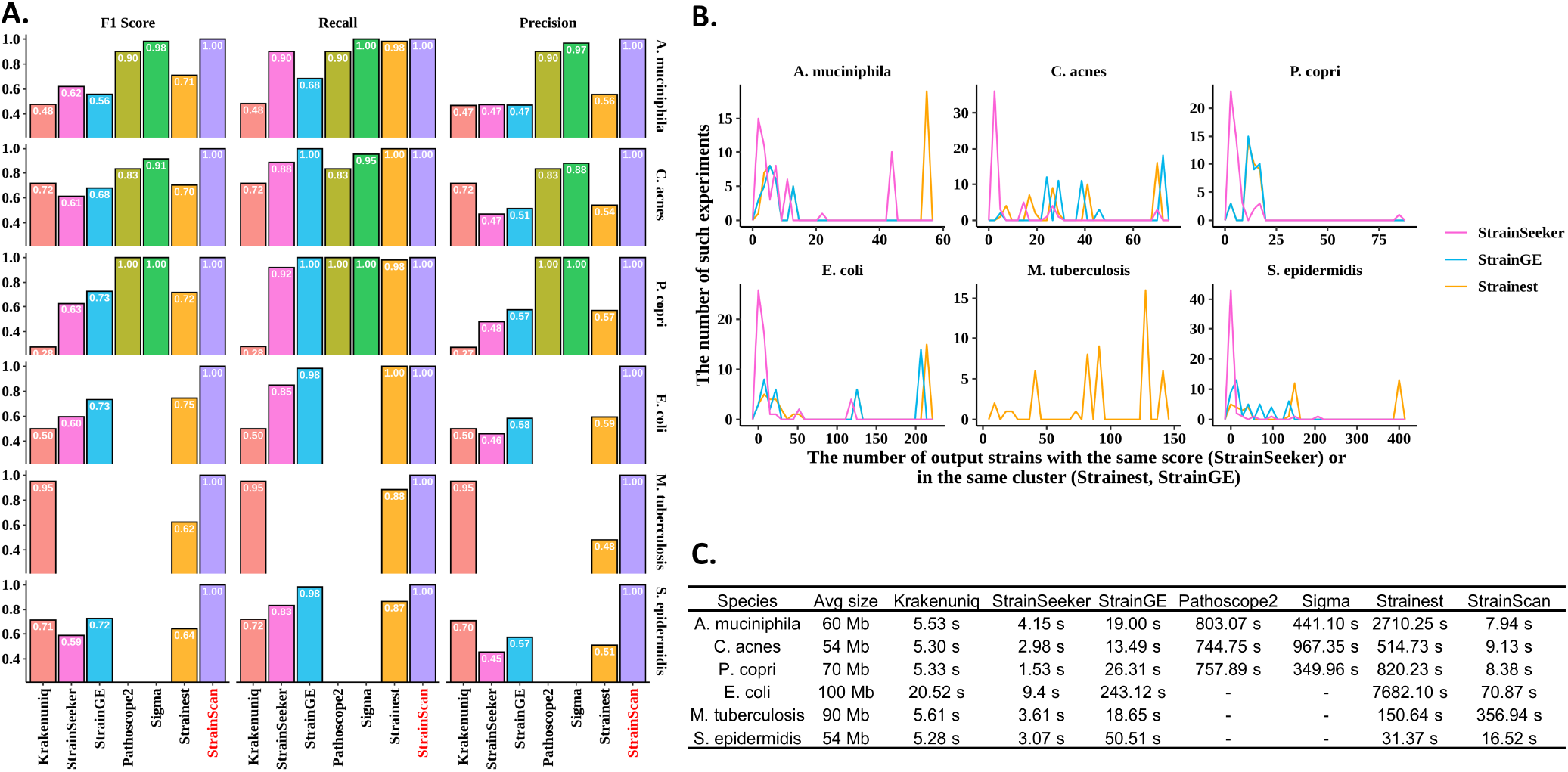
**(A).** The F1 score, recall, precision of 7 tools on “single-strain” simulated datasets. There are 60 sets of simulated reads for each bacterial species. **(B).** The histogram of the identical outputs of three tools. X-axis means the number of output strains with the same score (StrainSeeker) or in the same cluster (Strainest, StrainGE). Y-axis means the number of such experiments. For example, “50” on the X-axis means that the tool outputs 50 strains with the same score and one of them is correct. **(C).** Running time comparison of 7 tested tools. Sigma and Pathoscope2 are not shown on some datasets because they are too computationally expensive to construct databases for the corresponding bacteria or identify strains from simulated reads.

### Detecting co-existing strains from simulated data

It has been shown that the human-associated microbiota is often a complex mixture of closely related strains of the same species^11^. To quantitatively compare the performance of Krakenuniq, StrainSeeker, StrainGE, Strainest, and StrainScan on identifying multiple strains of the same species, we generated simulated datasets containing 2, 3, and 5 randomly selected strains from six bacterial species. Because Sigma and Pathoscope2 took a lengthy time to process these datasets, they were not included in this experiment.

To investigate how the similarities between the strains affect the tool’s performance, we used two strategies in the selection of multiple strains. During the clustering step of StrainScan, strains with *k*-mer-based Jaccard similarity greater than or equal to 95% (corresponding to an approximate ANI of 99.89%) are grouped to the same cluster. Therefore, it is more difficult to identify and distinguish the co-existing strains that are in the same cluster than those from different clusters. To consider different levels of difficulty, our first strategy randomly picked strains from different clusters while the second strategy selected different strains from the same cluster. For each strategy, we randomly selected 2, 3, and 5 strains (3 groups) and simulated the short reads using different coverage profiles: 100X and 10X for 2 strains, 100X, 50X, and 10X for 3 strains, and 100X, 70X, 50X, 20X, and 10X for 5 strains. Other read simulation parameters are the same as the “single-strain” experiment. Then we repeated the experiment 10 times by choosing another group of strains. Ultimately, for each bacterial species, we generated 30 sets of data containing different numbers of strains using the first and the second strategies, for a total of 60 sets of data. So there were a total of 360 (60×6) simulated datasets for the six bacterial species.

In order to compare the performance of different tools, we calculated the recall, precision, and F1 score. The performance comparison is shown in Figure 3A. StrainScan achieves near-perfect F1 scores on all tested datasets. Though StrainGE and Strainest have close recall to StrainScan in some bacterial species, their precision is low. Besides, Strainest relies on alignment, and therefore the identification time would increase quickly with the data size and data complexity (Figure 3C). For the two faster tools, Krakenuniq and StrainSeeker, neither performed very well. Among them, Krakenuniq performed better in identifying strains from different clusters than in identifying strains from the same cluster, which was in line with its method. StrainSeeker did not perform well in both cases and had worse performance in the datasets with higher complexity, indicating that it was unsuitable for identifying data with multiple strains. Overall, compared to other tested tools, StrainScan achieves more than 20% improvement in F1 score for datasets containing strains from the same cluster and has the highest F1 score in all tested datasets. Same as the “single-strain” experiment, we also investigated the number of strains falsely identified by StrainSeeker, StrainGE, and Strainest (Figure 3B). As shown in Figure 3B, there are more false positive hits in the right panel compared to the left panel, which suggests that the high similarity between strains leads to more false positive results for these tools. In addition, the high similarity also increases the number of strains that cannot be further distinguished (e.g. same score) in the experiment. For example, StrainGE and Strainest falsely identified more than 100 strains for *E. coli* strains from the same cluster but only falsely identified less than 15 strains for those from different clusters. Furthermore, there are a large number of SNVs between the actual strains and the representative strains in the databases of StrainGE and Strainest (see Supplementary Figure S5). Meanwhile, StrainScan has a more competitive speed compared to StrainGE and Strainest (Figure 3C). However, as mentioned in the last section, due to the high similarity between strains of *M. tuberculosis*, StrainScan sacrifices the computational efficiency to distinguish the strains in the same big cluster and thus took a long time to process datasets of *M. tuberculosis*. In conclusion, these results demonstrate that StrainScan performs better with complex samples than other tools, returning more accurate results even when there are a large number of similar strains in the sample with different sequencing depths.

**Figure 3.**
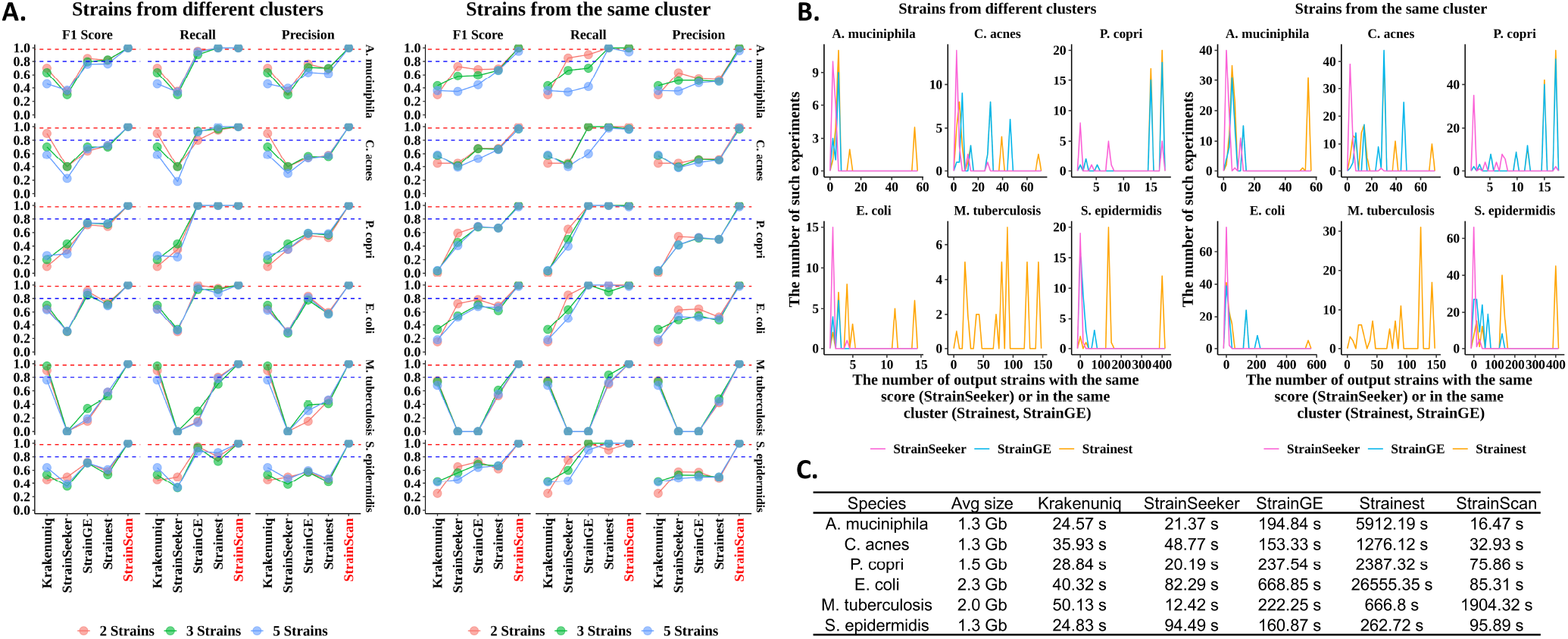
**(A).** The F1 score, recall, precision of 5 tools on “multiple-strain” simulated datasets. There are 60 sets of simulated reads containing 2, 3, and 5 strains with different similarities for each bacterial species. Note that StrainSeeker is not able to identify strains of *M. tuberculosis* and therefore the related scores are 0. **(B).** The histogram of the identical outputs of three tools. X-axis means the number of output strains with the same score (StrainSeeker) or in the same cluster (Strainest, StrainGE). Y-axis means the number of such experiments. Y-axis means the number of such experiments. For example, “50” on the X-axis means that the tool outputs 50 strains with the same score and one of them is correct. **(C).** Running time comparison of 5 tested tools.

#### Relative abundance computation

In order to measure the accuracy of the predicted strain profiles in synthetic data sets, we computed the Jensen–Shannon divergence (JSD) between the actual and the inferred frequencies. In case the dimension of predicted and true relative abundance may be different, we add zeros to the one with a lower dimension to calculate JSD. The result is shown in Figure 4. Figure 4 shows that in all cases the strain distribution reconstructed by StrainScan had high precision, with a median of Jensen–Shannon divergence (JSD) < 0.05. Although StrainGE and Strainest also had a good performance in most cases, they had low resolution and only returned one representative strain for the “Similar” datasets, which actually contained multiple strains. As a result, these two tools had worse performance in the “Similar” datasets, with a median of Jensen–Shannon divergence (JSD) > 0.05. For StrainSeeker and Krakenuniq, the JSD was bigger than 0.1 in most cases, indicating that these tools cannot accurately quantify the composition of strains. With increasing strain numbers and similarity, the median of JSD values of most tested tools increases while StrainScan’s JSD doesn’t fluctuate much in all cases. These results show that StrainScan can better quantify the composition of complex samples than other tools, even for samples containing highly similar strains.

**Figure 4.**
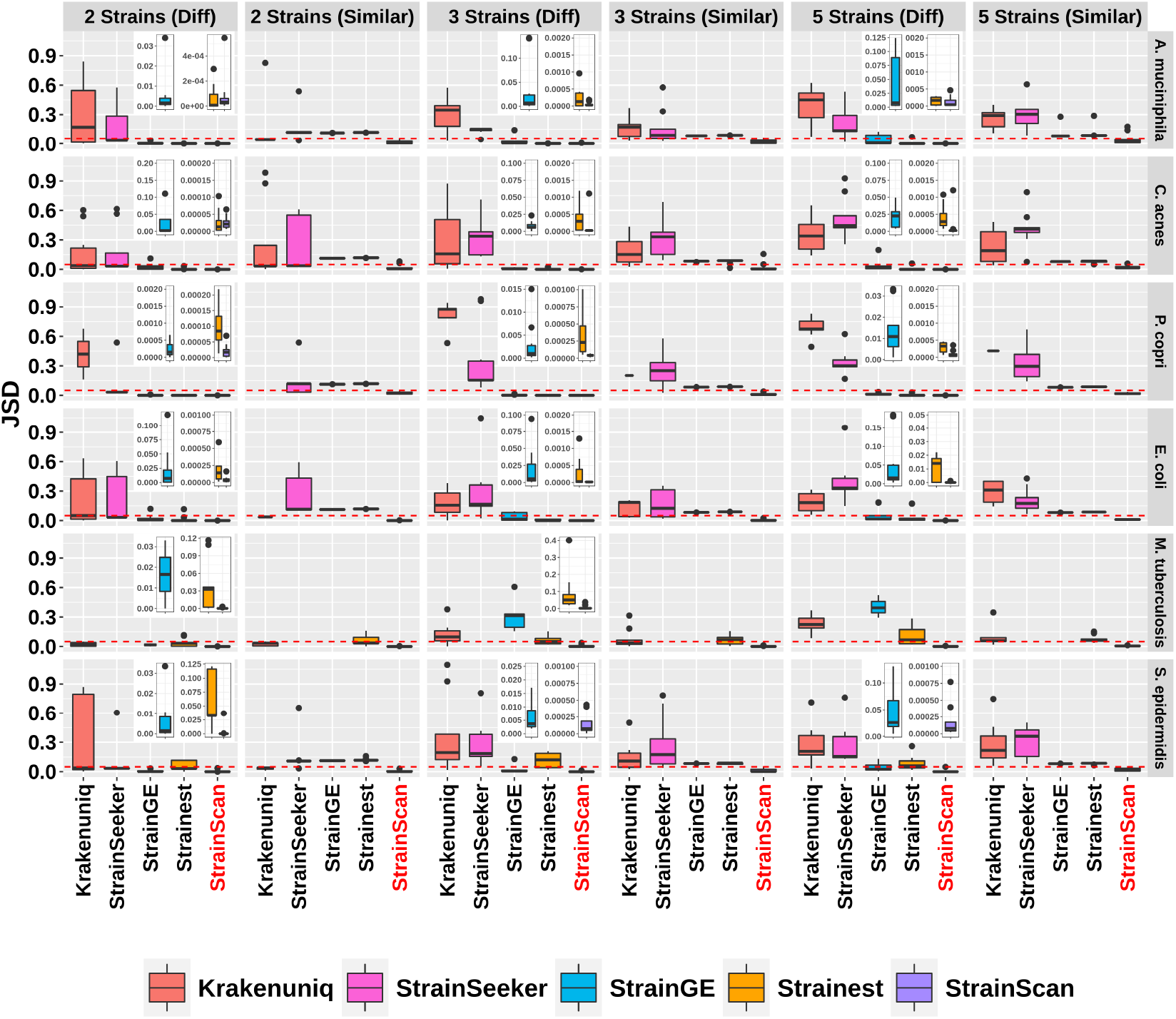
The Jensen-Shannon Divergence (JSD) of 5 tools between the ground truth and predicted relative abundance. “Diff” refers to strains from different clusters with *k*-mer Jaccard similarity < 95%, and “Similar” refers to strains from the same cluster with *k*-mer Jaccard similarity ≥ 95%. The red line in the figure refers to 0.05 of Jensen–Shannon divergence (JSD). The small windows in some of the plots show the JSD distribution for StrainGE, Strainest and StrainScan on a much smaller scale.

### Evaluation of StrainScan on the HMP mock data and *E. coli* mock community

#### The HMP mock data

In this experiment, we tested StrainScan on the two communities from the Human Microbiome Project^33^. They contain 21 known organisms with even (SRR172902) or staggered composition (SRR172903). Out of the 21 organisms, 3 bacteria (*E. coli, C. acnes*, and *S. epidermidis*) represent hard cases for strain-level analysis and we have established reference indexing structures. Thus, we tested StrainScan on strain-level analysis of the three bacteria. For the three bacteria, each one has only 1 strain in these two datasets. This test was especially challenging, due to the low abundance of some of these species in these samples. For comparison, we also used Krakenuniq, StrainSeeker, StrainGE, and Strainest to identify the strains of these bacterial species in these two datasets. The results of these 5 tools are shown in Table 1.

**Table 1.**
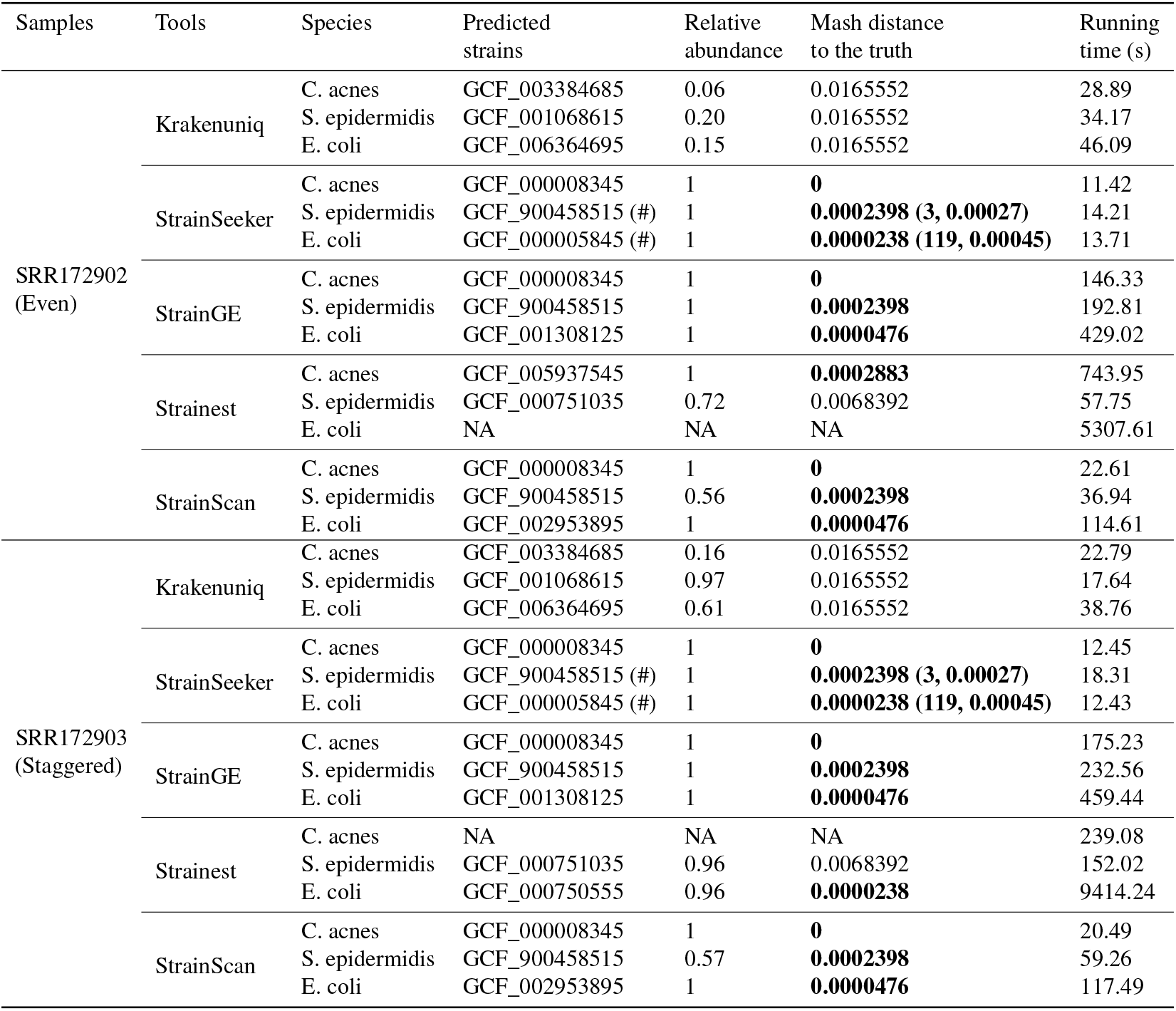
Analysis of two mock communities from the HMP project. Each bacteria has only one strain in the dataset, thus the relative abundance should be 1.0. For StrainSeeker, we took the strain with the smallest Mash distance to the truth as the predicted dominant strain. “(#)” means there are multiple hits with identical scores in the output. “NA” in the table indicates missing values. The character in bold means the Mash distance of predicted dominant strain to the truth is < 0.05%. For StrainSeeker, the two numbers in parentheses represent the number of multiple hits and the average mash distance between all hits and the ground truth, respectively.

For all tested species, StrainScan correctly identified the presence of one dominant strain that is highly similar (Mash distance to the truth < 0.05%) to the bona fide strain. Besides StrainScan, StrainSeeker and StrainGE also returned strains that are highly similar to the ground truth. However, the output of StrainSeeker often contains multiple hits with identical scores, making accurate evaluations difficult. For example, it returns 119 *E. coli* strains in two tested datasets, which makes it hard for users to know the actual strain present in these samples. In Table 1, we take the strain with the smallest Mash distance to the truth as the predicted dominant strain by StrainSeeker. StrainGE also returns strains with a small Mash distance to the ground truth, but these strains belong to clusters containing many other strains, which also makes it hard to know the actual strains present in the sample. Of the two remaining tools, Strainest was unable to identify low-abundance strains and it took a long time to run, while Krakenuniq returned results that differed significantly from the ground truth.

#### The E. coli mock community with multiple strains

To evaluate each tool’s performance on a real-world sample with a known multi-strain composition, we downloaded a mock community sequencing dataset (SRR13355226), which contains a large number of reads from the host (i.e. human) as well as four different *E. coli* strains. All the reads are used as input to all tools. StrainScan was the only tool that identified four strains correctly, with no false positive identifications (Figure 5). While StrainGE and Strainest correctly identified the same four strains, they returned representative strains that are not the ground truth. Furthermore, StrainGE identified a cluster containing 205 strains and Strainest identified strains that are not present in the mixture. The remaining two tools did not correctly identify all four strains, among which StrainSeeker’s output contained multiple hits with identical scores, while Krakenuniq reported many false positive strains.

**Figure 5.**
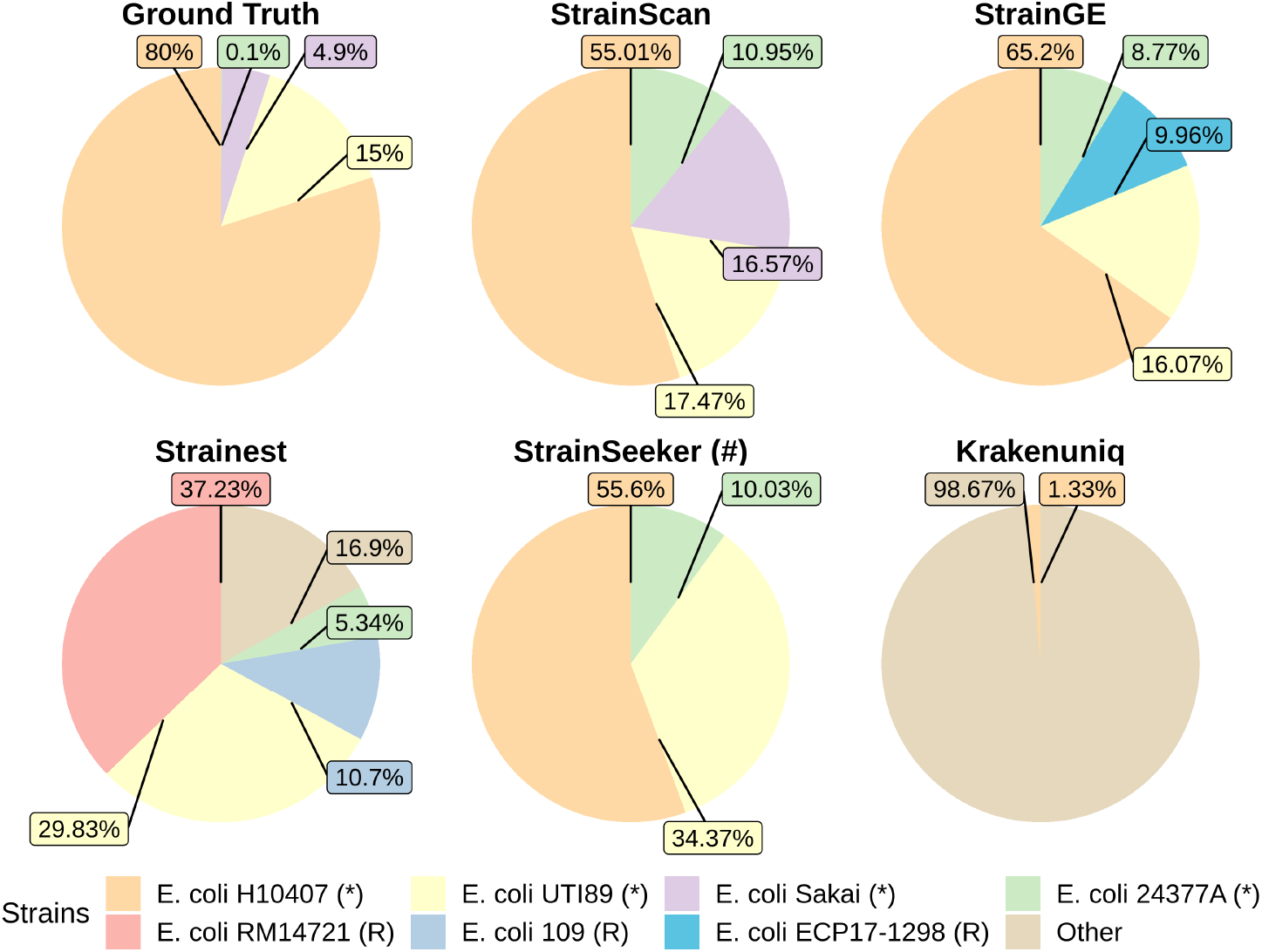
Analysis of the mock *E. coli* community data (SRR13355226). “(#)”: there are multiple hits with identical scores in the output. “(*)”: strains present in the ground truth. “(R)”: the identified strain is the representative strain of the ground truth. “RM14721” is the representative strain of “H10407” in Strainest. “Sakai” is the representative strain for “ECP17-1298” and “109” in StrainGE and Stainest, respectively.

### StrainScan detects the pathogenic strain from real sequencing data

To illustrate the potential application of StrainScan in pathogen detection, we have applied StrainScan to examine the presence of the pathogenic strain of *E. coli* and *M. tuberculosis* in two studies (BioProject Accession: PRJEB1775 and PRJEB2777). The first study is related to the 2011 Escherichia coli outbreak in Germany^34^. The outbreak was caused by an enteroaggregative (EAEC) strain O104:H4. The second study investigates the frequency of *M. tuberculosis* relapses within patients from the REMoxTB clinical trial, which evaluated the treatment for *M. tuberculosis* in previously untreated patients^35^. For each sample sequenced in this study, there is a known label representing the strain contained in that sample. From each of these two studies, we selected six samples for the experiment. For comparison, we also applied other 4 computationally efficient tools to detect the pathogenic strains in these real sequencing data. As shown in Table 2, StrainScan was able to identify the correct strains in all tested datasets while other tools failed to detect correct strains in some datasets. Although Strainest could identify correct strains in most datasets, it only returned the representative strain of the correct strain for some datasets.

**Table 2.** Comparative performance of StrainScan, StrainGE, KrakenUniq, Strainest, and StrainSeeker for identification of pathogenic strain of *E. coli* and *M. tuberculosis*. The results marked “Y” in the green block mean these results are consistent with the ground truth while “N” in the red block indicates the inconsistent result. “(#)” means there are multiple hits with identical scores in the output and “(R)” means the identified strain is the representative strain of the ground truth. Note that StrainSeeker is not able to identify strains of *M. tuberculosis* and therefore related results are “-”. For StrainSeeker, the number in parentheses represent the number of multiple hits.

| Species | SRA accession | Known strain | StrainScan result | StrainGE result | Krakenuniq result | Strainest result | StrainSeeker result |
| --- | --- | --- | --- | --- | --- | --- | --- |
| *E. coli* | ERR260487 | O104:H4 | Y | Y | N | Y | Y (#, 5) |
|  | ERR260492 | O104:H4 | Y | Y | N | Y | Y (#, 5) |
|  | ERR260499 | O104:H4 | Y | Y | N | N | Y (#, 24) |
|  | ERR260491 | O104:H4 | Y | Y | N | Y | Y (#, 5) |
|  | ERR260480 | O104:H4 | Y | Y | N | Y | Y (#, 5) |
|  | ERR262949 | O104:H4 | Y | Y | N | Y | Y (#, 5) |
| *M. tuberculosis* | ERR751569 | GCF_900127125 | Y | N | N | Y (R) | - |
|  | ERR751369 | GCF_900120835 | Y | N | Y | Y | - |
|  | ERR751411 | GCF_900120975 | Y | N | Y | N | - |
|  | ERR751393 | GCF_900120575 | Y | N | N | Y (R) | - |
|  | ERR751396 | GCF_900120785 | Y | N | Y | Y (R) | - |
|  | ERR751382 | GCF_900120755 | Y | N | Y | Y (R) | - |

### StrainScan reveals the greater diversity of *C. acnes* in human skin

A recent study11 shows that *C. acnes* is one of the most common bacteria on human skin and usually has a complex multi-strain community. Strainest was applied to re-analyzed the human skin data set (SRP002480) from the study^20^. However, we found that in some samples, the number of strains identified by Strainest was often less than the number of strains reported by the original study. In the original study, the authors used an in-house pipeline to determine strains in the samples, and the accuracy of this in-house pipeline was previously validated with extensive simulations for human skin microbiome data^10^. Therefore, we selected nine samples from two individuals who fit this case and then re-analyzed these samples using Strainest and StrainScan. Then, we compared the predicted results with those reported in the original study. For consistency, we selected all strains used in the original study to build the new custom databases for Strainest and StrainScan, and used these newly established databases for subsequent analyses. The result is shown in Figure 6. Strainest only returned one or two strains in all tested datasets while more strains are reported according to the original study. Meanwhile, StrainScan displayed a more similar relative abundance pattern with the reported result than Strainest. Besides, StrainScan also detected some strains in the samples of “HV05_AI” that were not found in the original study, which implied the greater diversity of *C. acnes*. For example, the strain “GCF_000145115” detected by StrainScan in the first sample of “HV05_AI” was highly similar to the strain “GCF_004631215” detected by Strainest in the same sample (Figure 6B). However, these strains were not reported in the original study.

**Figure 6.**
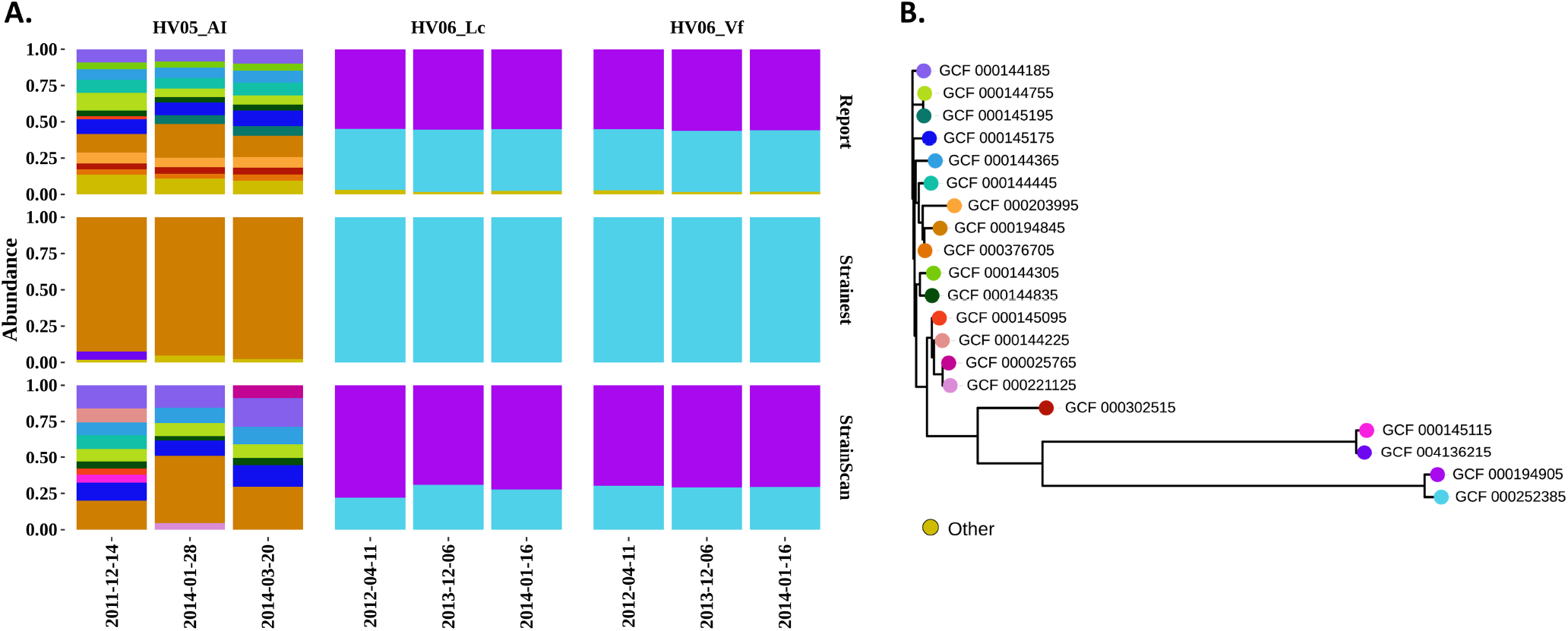
(A) StrainScan reveals the greater diversity of *C. acnes* in 9 real metagenomic samples. These samples were taken from three different sites of the skin of two healthy individuals, at different time points. The site codes are described in the original work^11^. (B) Phylogenetic tree of the identified strains. Leaves are colored using the same schema as in (A) and the distance is the Mash^36^ distance.

### Analysis of real sequencing data in cross-sectional studies

Two studies^16, 20^ have demonstrated a correlation between the distribution of *E. coli* strains and the geographical location of their hosts. StrainScan was applied to derive the correlation using datasets from different studies. To show this utility, we applied StrainScan to determine the strain distribution of *E. coli* in several recent studies. We have analyzed two sets of metagenomic and one set of whole genomic sequencing sample from stool or blood samples of different studies, with one dataset including 6 infants in Estonia^37^, the second including 6 Chinese adults^38^, and the third including 6 USA adults (PRJNA278886). Considering that these samples are from different studies and countries, we only keep the dominant strain of each analyzed sample. As a result, StrainScan distinguishes *E. coli* strains into three distinct groups (Figure 7A). According to the phylogenetic tree analysis using Prokka^39^ and Roary^40^, these identified strains also fall into three distinct clades (Figure 7B).

**Figure 7.**
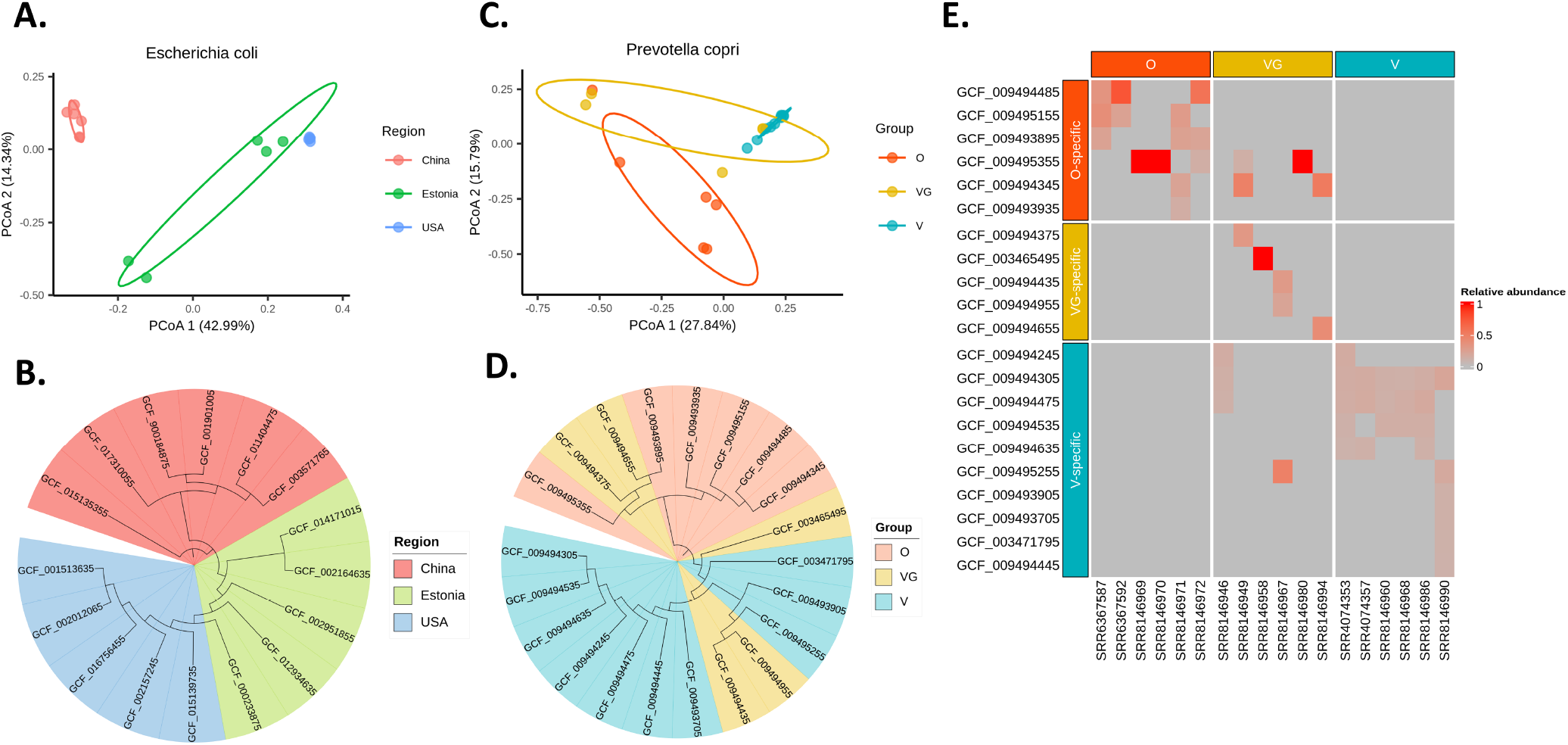
StrainScan profiling of *E. coli* and *P. copri* from metagenomics samples. (A-B) *E. coli* genomic diversity in the healthy sample of Chinese, Estonian and American cohorts as seen by principal coordinates analysis (PCoA) and the phylogenetic tree analysis of dominant strains. (C-D) *P. copri* genomic diversity in the healthy gut of Omnivores (O), Vegetarians (VG), and Vegans (V) cohorts as seen by principal coordinates analysis (PCoA) and the phylogenetic tree analysis of specific strains. (E) Strain relative abundance profiles of the 21 identified strains in the 18 metagenomic samples from three groups.

### Investigation of strain diversity of *P. corpi* in different populations

The evolution of some host-associated microbes can be linked to factors such as host diet. For example, one recent study shows that the host diet and the distinct genetics of human intestinal *P. copri* strains are closely related^8^. However, the original study mainly explored the diversity of strains within different populations from a functional perspective, and not from a taxonomic perspective. Thus, we applied StrainScan to re-analyze 18 human gut metagenomic datasets that are from the same study and have high abundance of *P. corpi*. As mentioned earlier, these samples were all taken from the study investigating diet and strain diversity^8^. According to the original study, these samples can be divided into three groups: Omnivores (O), Vegetarians (VG) and Vegans (V). We selected six samples from each group, all six of which had high abundance of *P. copri* strains. Then, we applied StrainScan to these samples that contain multiple strains of *P. copri* and selected the strains with relative abundance > 0.02 for further analysis. The relative abundance profile of each sample is shown in Figure 7E. In Figure 7E, we can find some strains are specific to one group, and therefore we label these strains as “specific” strains of that group. These “specific” strains are then analyzed using the principal coordinates analysis (PCoA) (Figure 7C) and the phylogenetic tree analysis (Figure 7D). The analysis result shows that “V-specific” strains are clearly separated from “O-specific” strains. Interestingly, “VG-specific” strains lie somewhere in between, which is consistent with the conclusion of the original study and our general knowledge.

## Discussion

In this work, we presented StrainScan, a new strain-level composition analysis tool for short reads. We designed a novel tree-based *k*-mer indexing structure to strike a balance between the strain identification accuracy and the computational complexity. Then, by applying informative *k*-mers and the elastic net model to identify strains and predict their abundance, StrainScan improved the resolution of the strain-level analysis and the accuracy of abundance estimation.

StrainScan shows higher accuracy and resolution than other tested tools across all benchmark datasets with different complexity. Especially, StrainScan outperforms all other tools on datasets containing strains with higher similarity and varied sequencing depth. This level of high resolution can be achieved by alignment-based tools such as Pathoscope2 and Sigma. But StrainScan is at least 10 times faster than them. The experiment results of mock data and real data further demonstrate that StrainScan has the potential for a more comprehensive strain-level composition analysis.

We tested the limits of StrainScan on building the CST for 25,349 complete and draft *E. coli* genomes. The program requires > 1TB memory. Our empirical tests show that StrainScan is efficient for building the CST for less than 5000 genomes. Thus, we recommend users to use only complete genomes for constructing the indexing structure if there are more than 5000 genomes. Our first step based on cluster search can efficiently reduce the search space. However, if all the reference genomes are highly similar and only differ by a handful of bases, they tend to be grouped in one cluster. In this case, the cluster search still returns a large search space for the second step, which does not take full advantage of the cluster indexing structure. *M. tuberculosis* has a large cluster, which slows down the strain identification.

For future work, we plan to extend some efficient data structures such as HyperLogLog^21^ to speed up both the database construction and identification process of StrainScan. Besides, we also plan to make StrainScan output more useful features such as SNVs and SVs between strains identified in the samples, which can be used to explore the function of different strains.

## Methods

### Overview of StrainScan

StrainScan is designed to identify known strains from short reads directly. Because there are many species-level composition analysis tools for metagenomic data, the inputs to StrainScan are the short reads in “fastq” format and strain genomes for targeted bacteria in “fasta” format. There are two major challenges for strain-level analysis. First, for highly similar strains, it is difficult to find sequence features (e.g. unique *k*-mers or SNVs) to distinguish them, resulting in lower accuracy and resolution. For example, Krakenuniq^21^ and StrainSeeker^22^, two strain identification tools utilizing *k*-mer-based features, performed poorly in distinguishing highly similar strains (the right panel of Figure 3A). Secondly, the large reference strain database poses a computational challenge for many alignment-based tools like Sigma^29^, Pathoscope2^30^, Strainest^20^, etc (Figure 2C and 3C). To strike a balance between the strain identification resolution and computational cost, we design a hierarchical indexing method that combines a fast but coarse-grained cluster search tree (CST) and a slower but fine-grained strain identification strategy inside a cluster. As shown in the flowchart in Figure 8, we will first create a cluster tree-based indexing structure. With our efficient and accurate cluster search method on this tree, we can first pinpoint a cluster that is present in the sample. Then we will use carefully chosen *k*-mers to distinguish different strains in the identified cluster or clusters. The hierarchical method has several advantages. First, it allows us to accommodate the heterogeneous similarity distribution between strains with some strains sharing much higher similarities than others. The strains with low similarity can be quickly identified by our fast CST search strategy. And only those highly similar strains need to use a more careful strategy in the second step. Second, the hierarchical method can increase the search accuracy by allowing us to use more unique *k*-mers (Supplementary Table S1). Any *k*-mer that is shared between clusters now can be utilized for within-cluster search. Third, the hierarchical method can reduce the member footprint. Without the hierarchical method, we need to search strains from all reference sets that contain a large number of *k*-mers. Given the clusters identified by CST search, StrainScan only needs to search strains in identified clusters that contain fewer strains and *k*-mers. For example, the total number of *k*-mers in *E. coli* reference set before clustering is 192,325,016, while the number of *k*-mers in the largest cluster after clustering is 16,071,080, more than a tenfold reduction (Supplementary Table S1).

**Figure 8.**
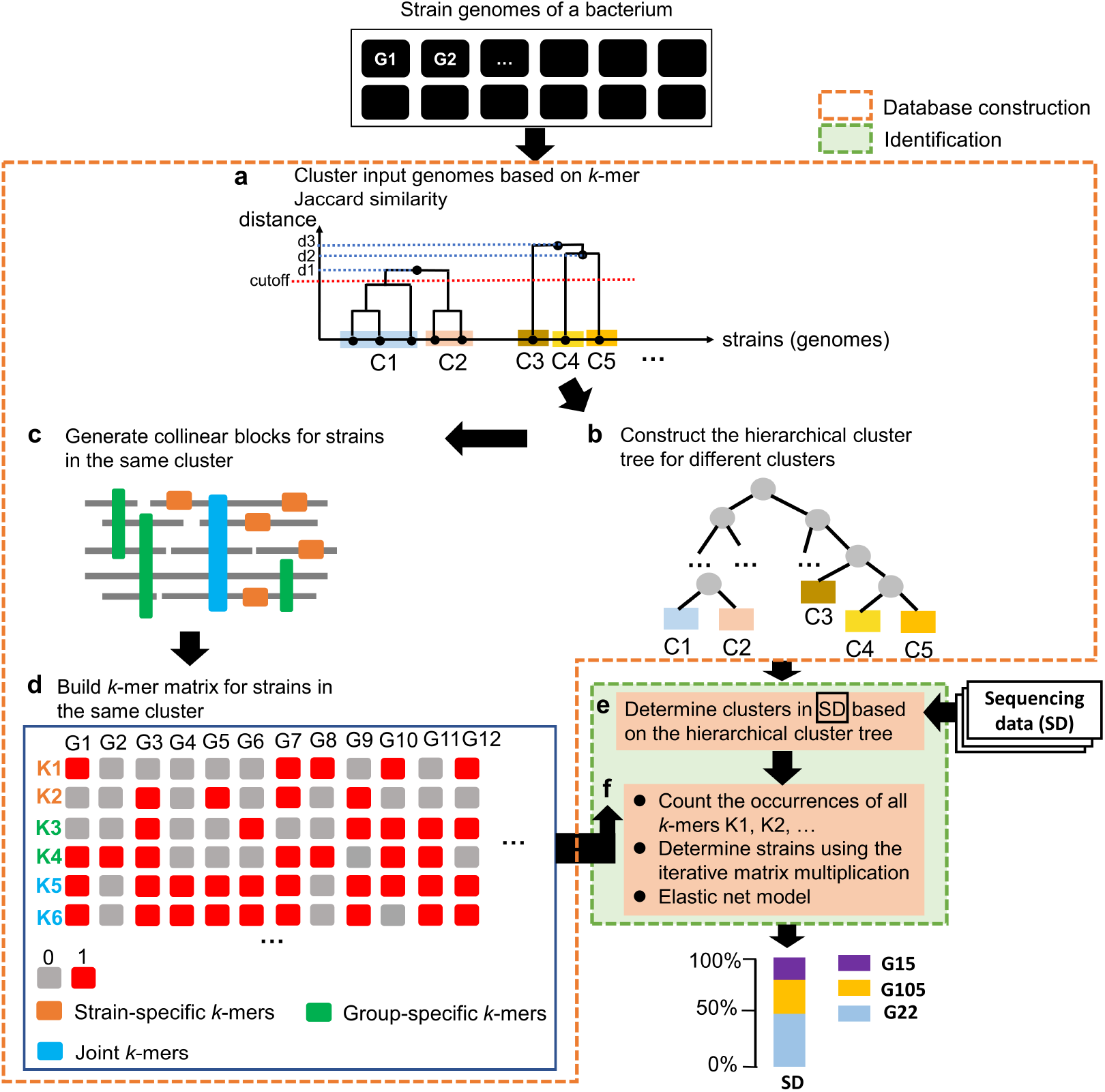
The overview of StrainScan. (a). The sketch of the strain genome clustering process. Given the strain genomes (G1, G2,…) of the bacteria of interest, all-against-all *k*-mer Jaccard similarities are computed using Dashing^31^. Genomes are then clustered using single-linkage hierarchical clustering. By default, the clustering threshold is set to a Jaccard similarity of 0.95. In this example, given the cutoff represented by the dashed red line, five clusters from C1 to C5 are output by the clustering process. (b). Given the clusters, construct the hierarchical cluster tree for later cluster-level identification. (c). Generate collinear blocks to extract *k*-mers that can help distinguish different strains inside the same cluster. (d). Step d concludes the indexing structure process for the reference genomes. (e and f). The indexing structure and the sequencing data (reads) are input for strain search. e: search for clusters. f: strains are identified by the iterative matrix multiplication and the relative abundance profile is finally inferred by elastic net regression.

### Construct the cluster search tree (CST) from highly similar strains

Given many strains’ genomes of the same species, we first calculated a Jaccard similarity matrix with an alignment-free, *k*-mer based method Dashing^31^ (*k* = 31). Then, we performed the agglomerative hierarchical clustering (single-linkage) based on this matrix, grouping the strains into a dendrogram. Finally, we chose a fixed height cutoff *H* (0.95 by default) to cut the dendrogram into many clusters, each of which consists of one or more strains. The strains inside each cluster have the *k*-mer-based Jaccard similarity ≥ 0.95, which roughly corresponds to average nucleotide identity (ANI) of 99.89%^19^.

To pinpoint the cluster where a strain is contained, we will convert the clusters and the dendrogram into a cluster search tree (CST) to support both accurate and efficient cluster search. The CST keeps the same tree topology as the dendrogram except that each cluster is represented by a leaf node in the tree. In addition, we discard the distance information in the dendrogram so that the distance between each node and its parent (or child) is uniform, regardless of their Jaccard similarity. Thus, the CST is a full binary tree. In order to support the cluster search, each node contains a set of *k*-mers that are unique to the subtree rooted by this node. By conducting *k*-mer match, the CST will guide us to take either the left child or the right child until reaching one or multiple leaf nodes (i.e., clusters). We first describe how we assign *k*-mers for each node.

### Construct the CST-based indexing structure

#### *k*-mer assignment for the nodes

A CST is defined by two elements: the tree topology and the *k*-mer set assigned for each node. In this section, we will describe how we assign *k*-mers to the nodes to support the cluster search. For a node *v* in the CST, we denote the subtree rooted by *v* as *v^T^*. The *k*-mer assignment for v follows two criteria. First, the *k*-mers should be shared by most of the strains in the leaf nodes of *v^T^*. Second, the *k*-mers are unique to the strains in *v^T^*. The two criteria are visualized using an example in Figure 9A.

**Figure 9.**
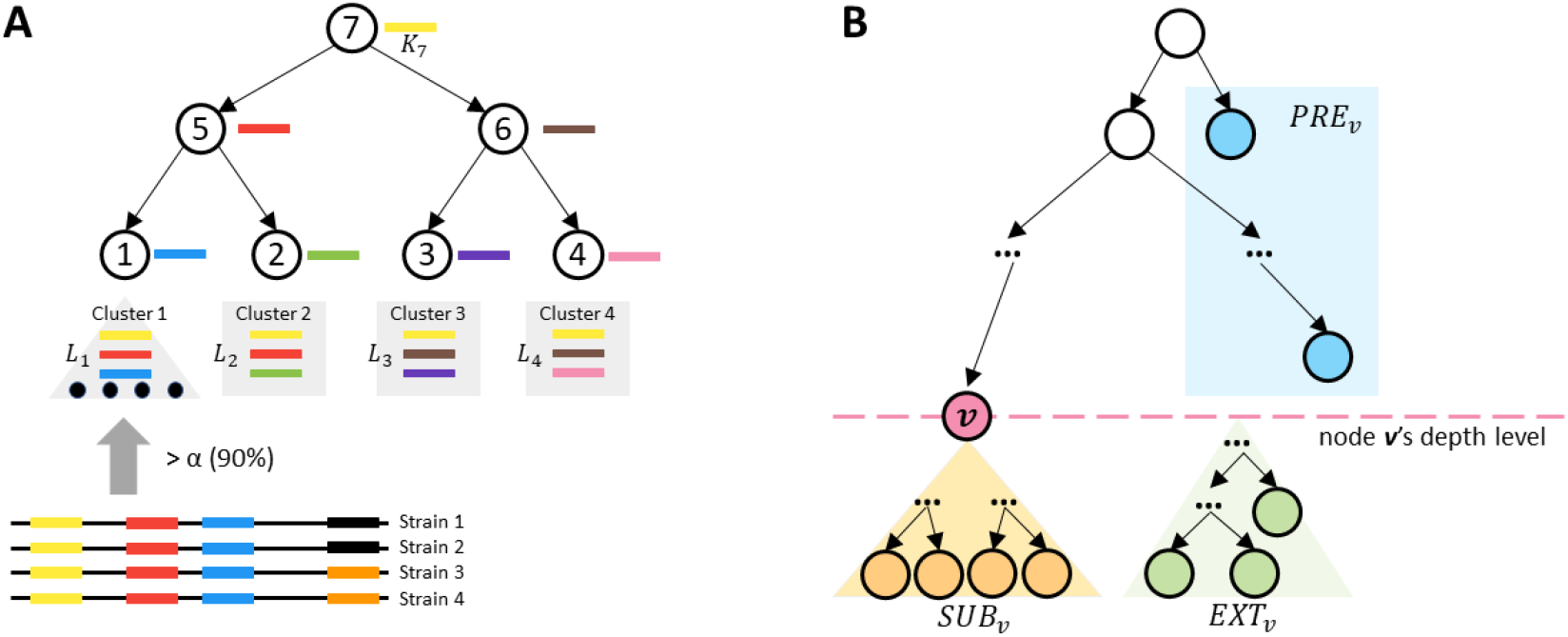
(A) An example of the *k*-mer assignment in the CST-based indexing structure. Each node possesses *k*-mers unique to its rooted subtree and shared by most of the strains in the subtree. Each bar with a specific color represents a *k*-mer and each node is assigned with one unique *k*-mer in this example. (B) When constructing node *v*’s *k*-mer set, all leaf nodes will be divided into three groups named *PRE_v_*, *SUB_v_*, and *EXT_v_*.

Following the two criteria, we first assign leaf nodes with *k*-mers extracted from strains in their corresponding clusters. To use *k*-mers that represent relatively well-conserved features in the underlying strains, only the *k*-mers that appear in at least α% of the strains will be kept for clusters with multiple strains. Big α indicates that only *k*-mers shared by many strains are used for building the CST while small α allows the CST to use strain(s)-specific *k*-mers. We compared the cluster identification performance using a range of α in our experiments. According to the empirical results in Supplementary Figure S1, we set the default α as 90.

Hereafter, we denote the initial *k*-mer set for a leaf node v as **L_v_**. Next, starting from the leaf nodes, we recursively move the shared *k*-mers between every two sibling nodes towards their parent. In the last step, all *k*-mers that occur in more than one node will be removed. At the end of this process, each node v (an internal node or a leaf node) contains a set of unique *k*-mers denoted as **K_v_**. Specifically, *K_v_* for a node v can be constructed using a set operation as shown in Equation (1). For the node v with depth *d_v_*, all the leaf nodes are divided into three groups based on their relationship with *v*, as shown in Figure 9B and defined in the box below.

***SUB_v_***: Leaf nodes in *v^T^*.
***EXT_v_***: Leaf nodes outside *v^T^* and with depths *d* ≥ *d_v_*.
***PRE_v_***: Leaf nodes outside *v^T^* and with depths *d* < *d_v_*.

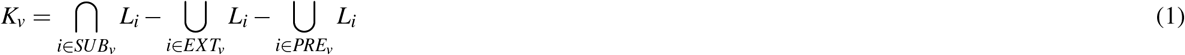

The CST constructed so far is similar to the tree built in StrainSeeker^22^. Although using the unique *k*-mers **K_v_** can guide the search for identifying strain clusters, a significant limitation is that some nodes only contain a small number of unique *k*-mers, which can lead to false positive (FP) matches more easily than nodes with many *k*-mers. This was observed when applying StrainSeeker. Take the StrainSeeker database built from 112 P. copri strains as an example. Out of 222 nodes, 21 nodes are empty, and 104 nodes have *k*-mers fewer than 1,000. The nodes with small *k*-mer sets tend to be matched by chance and thus lead to FP identification. In order to address this limitation, we will augment those nodes by adding *k*-mers that do not add ambiguity to the cluster search.

#### Optimize the CST

The main task of the optimization process is to reconstruct all small *k*-mer sets. In the CST, if a node has a large *k*-mer set, it is called a *strong node*. And the nodes with small *k*-mer sets are called *weak nodes*. To distinguish strong and weak nodes, we need to set a specific cutoff of the *k*-mer number.

##### Determination of the *k*-mer number cutoff

Ideally, we assume that reads are randomly distributed across a genome. Thus, the count of each *k*-mer follows a Poisson distribution with mean λ equal to the average coverage across the whole genome^41^. When a node only possesses a few *k*-mers, its *k*-mer count distribution will not reflect the coverage of the strains in a sample. This not only causes a high FP rate but also reduces the accuracy of strain abundance estimation. Thus, we require that each node has enough *k*-mers so that its *k*-mer counts still exhibit a Poisson distribution with a mean roughly equal to λ. If these *k*-mers are independently distributed, estimating the minimum *k*-mer number cutoff of the reliable *k*-mer set would be easy. Figure 10 shows how the *k*-mer set size influences the Kolmogorov–Smirnov (KS) distance between *k*-mer count distributions of a given *k*-mer set and the whole genome. When the *k*-mer number reaches 1,000, the KS distance decreases very slowly and approaches 0.02. Therefore, we set the *k*-mer number cutoff as 1,000 in this work.

**Figure 10.**
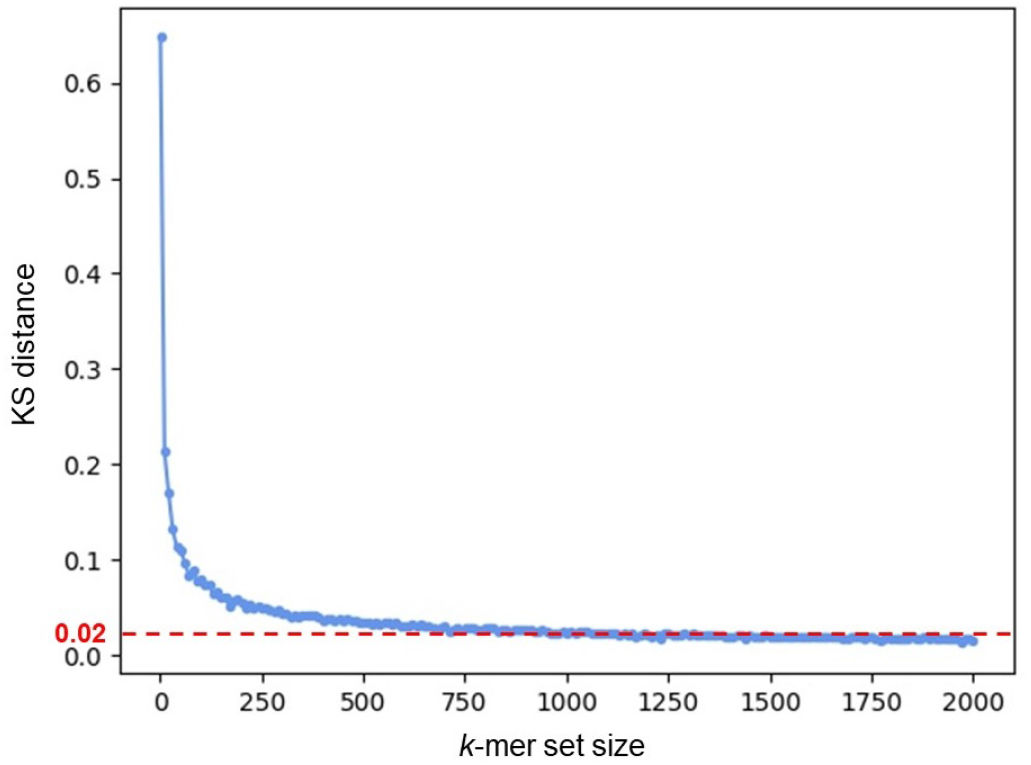
The change of the KS distance between the *k*-mer count distributions on a *k*-mer set and the *k*-mers of the whole genome with the *k*-mer set size. The *k*-mer set’s sizes range from 1 to 2,000 (X-axis). The *k*-mer set is randomly sampled from the genome multiple times. And the KS distances are the average values from these repeated experiments. The coverage of the simulated reads is 10x. When the KS distance is less than 0.02, there is no significant difference between the two distributions. where 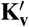 is the reconstructed *k*-mer set for the given weak node v. Compared with equation (1), a part of *k*-mers shared between *PRE_v_* and *SUB_v_* can be added to v’s *k*-mer set, thereby achieving the weak node augmentation.

Once we determine the cutoff, we can identify all weak nodes with *k*-mer numbers below the cutoff 1,000. A key observation behind the weak node augmentation is that we adopt Breadth-first Search (BFS) for the CST traversal and thus only need to choose among nodes at the same depth. As long as we can ensure that the nodes at the same depth have no overlapping *k*-mers, we can conduct the search without any ambiguity. This means that we can tolerate some non-unique *k*-mers in the nodes of CST. When we traverse to a node *v* at depth *d*, any leaf node *i* at depth smaller than *d* has been examined, and thus *k*-mers from *Li* might be reused by node *v*. If *v* is a weak node, we can change it into a strong node using this method. Specifically, we employ the set operations of *k*-mers to reconstruct every weak node:

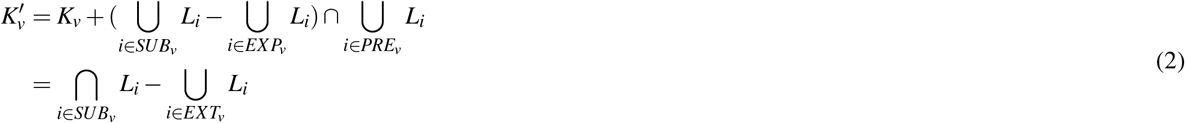

After implementing the reconstruction method, most of the weak nodes will become strong nodes. For instance, in the optimized CST built from 112 P. copri strains, only three out of 101 nodes are weak using our method. In order to decide the presence of a strain in a sample with high precision, we require all leaf nodes to be strong nodes. If the above procedure still leaves weak leaf nodes, we will combine them with their sibling nodes so that enough *k*-mers can be derived from the combined node. This process will iterate until all leaf nodes become strong nodes.

### Cluster search in the CST

Given the input sequencing data, we first extract all *k*-mers from the CST and conduct fast *k*-mer match for all short reads using Jellyfish^42^. Then, the *k*-mer match counts by all reads will be mapped back to the CST. Each node *v* will be assigned with a one-dimensional numerical vector **C_v_** = (*c*_1_,*c*_2_,…, *c*_|**Kv**|_), with each cell recording a *k*-mer match count. The cluster search algorithm is based on BFS, starting from the root and examining *k*-mer matches for nodes level by level (Figure 11). The *k*-mer match vector *C_v_* of each node v is used to decide whether or not to traverse v’s descendants based on a binomial test. The final search results contain one or multiple leaf nodes representing the strain clusters present in the sequencing data.

**Figure 11.**
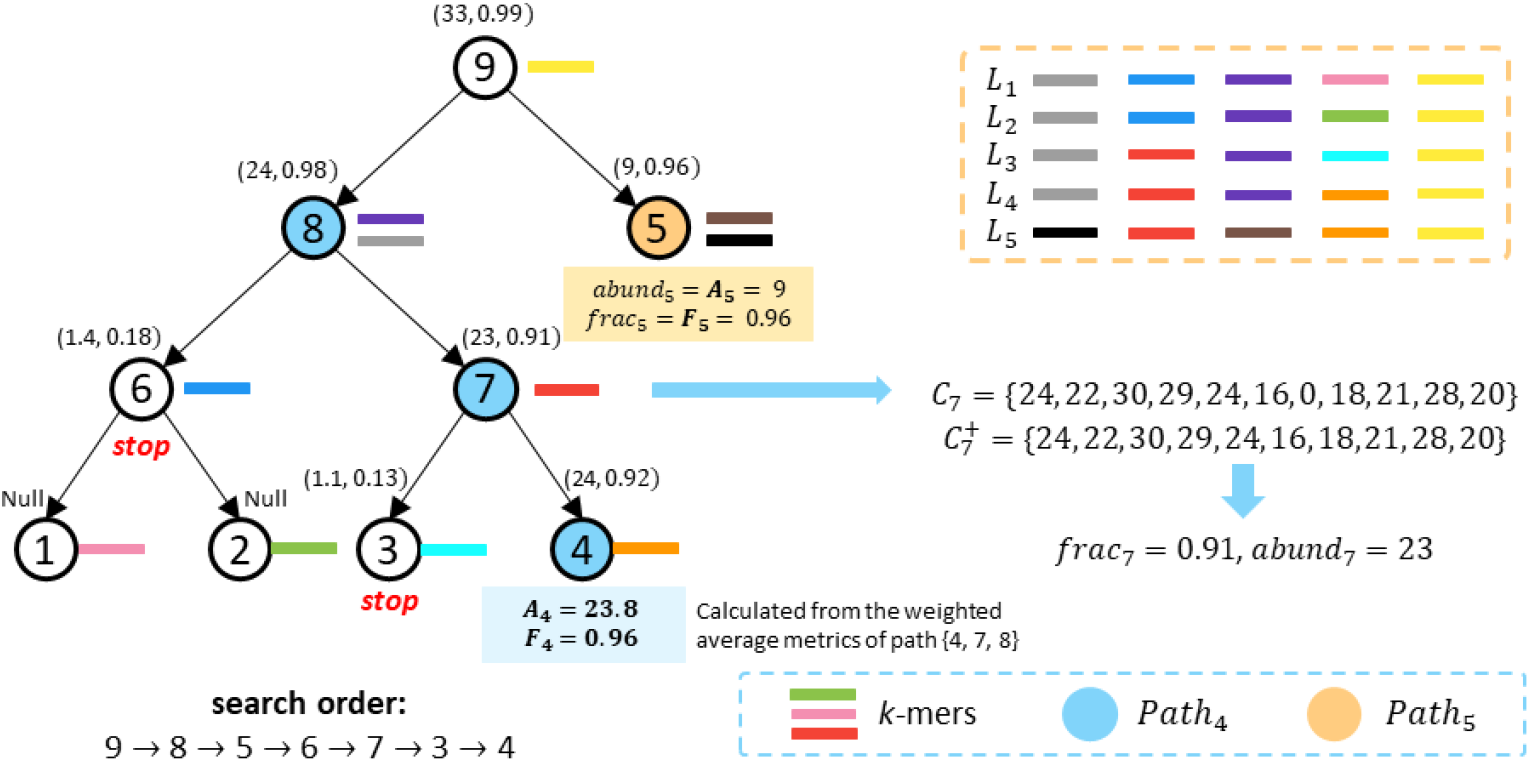
An example of the cluster search process. The values of the two scoring metrics (*abund_i_*, *frac_i_*) are shown beside each node *v*. The search results contain cluster 4 and 5 with estimated abundance 23.8 and 9, respectively. Node 6 and 3 did not pass the binomial test, thereby failing to traverse their descendants. The nodes in *Path*_4_ and *Path*_5_ are colored by blue and orange, respectively. *K_4_* shares the same orange *k*-mer with *L*_5_. Thus, we need to adjust *C*_4_ based on cluster 5’s estimated abundance *A*_5_ to calculate the accurate scoring metrics.

#### Scoring metrics for the cluster search

When the search visits a node v, two scoring metrics will be calculated to decide which child nodes will be visited. As shown in Figure 11, the first metric is the fraction of matching *k*-mers (**frac_v_**), which represents the fraction of *k*-mers in *K_v_* that is present in the sequencing data. It is defined as:

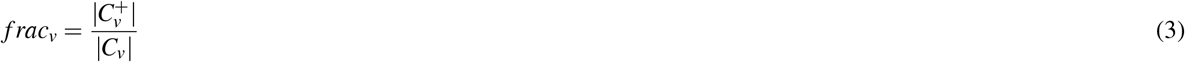

where 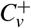 represents the vector which contains all positive *k*-mer counts from *C_v_*.

The second metric is the average *k*-mer match count (**abund_v_**), which is computed only using *k*-mers with positive matching counts. And when *frac_v_* < 0.1, *abund_v_* will be set to 0.

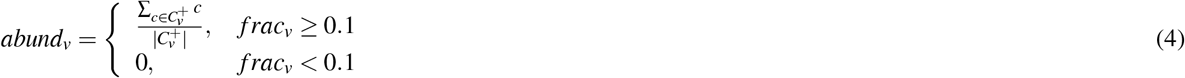

#### The search strategy for CST

After we calculate the two scoring metrics of v, we conduct a binomial test to decide the traversal order in CST^13^. Because sequencing errors can incur *k*-mer matches, the main goal of the binomial test is to distinguish random matches by sequencing errors from true matches for a true strain, which is particularly important for strains of low abundance. Given a node *v*, we examine whether we can reject the null hypothesis that *abund_v_* is generated by sequencing errors.

Specifically, we first round *abund_v_* and *abund_p_* (*p* is the parent node of *v*) to their nearest integers 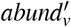 and 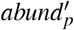. Then, given the sequencing error rate *e* (1% by default), we reject the error-caused null hypothesis when the probability of *abund_v_* being generated from sequencing errors is smaller than *β* (0.05 by default). The probability is estimated with

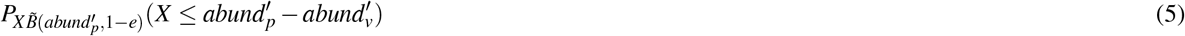

where *B*(*abund_p_*, 1 – e) is the probability mass function of the binomial distribution with 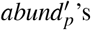 trials and the successful rate 1 – e. Failing to reject the null hypothesis indicates that we cannot distinguish low-coverage *k*-mer matches and sequencing noise. Thus, we consider *abund_v_* is just from sequencing errors, and the search will stop for *v*’s descendants. Otherwise, if we succeed in rejecting the null hypothesis, we believe that one or multiple strain clusters in *v^T^* are present in the sequencing data. And the CST search will add *v*’s two child nodes to the end of the BFS queue, preparing to traverse them later. Unlike the traditional binary search tree (BST), the two sibling nodes of the same parent can both reject the error-caused null hypothesis. Therefore, we can traverse all of their descendants, and the search results of CST are probably more than one.

#### Determination of the identified clusters

Once we reach a leaf node, we will further examine the *k*-mer matching statistics using both the leaf node and its ancestor nodes that contain *k*-mers moved from the leaf nodes. If there is only a single leaf node identified, all the nodes along the path from the root to *v* can be used to compute the *k*-mer statistics. However, if there are multiple leaf nodes identified, not all the ancestor nodes should be used. Instead, only the ones that contribute uniquely to the leaf node v should be used to compute the final abundance. These nodes can collectively constitute a path *Path_v_*, where all of the *k*-mer matches on *Path_v_* are only originated from strains in the leaf node v. To identify *Path_v_*, we first identify the maximum subtree that only contains v as the identified node. And *Path_v_* is equivalent to the path between the root of this subtree to v. Using the *k*-mer counts of nodes on *Path_v_* all together to estimate the cluster’s abundance will provide higher confidence than using a single leaf node. Take *Path*_4_ in Figure 11 as an example, two leaf nodes 4 and 5 are identified. In this case, *Path*_4_ is the root-to-leaf path containing *v* in node 8’s rooted subtree 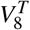. Subsequently, we can collect all *k*-mer match counts *C_i_* on *Path_v_* to calculate the weighted average fraction of matching *k*-mers **F_v_** and the weighted average *k*-mer match count **A_v_**:

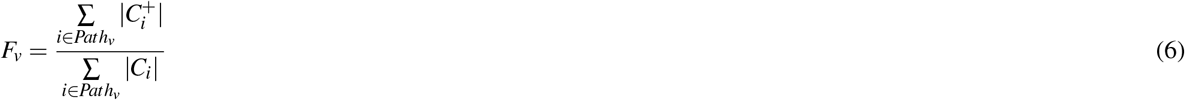

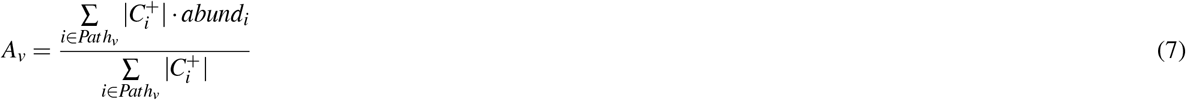

If *F_v_* is larger than a given cutoff (the default value is 0.4, but users can modify the value to adapt to different conditions), we consider the cluster in v is present in the sequencing data. After finishing the CST search, all identified clusters will be output with their estimated abundances (calculated by *A_v_*).

#### Adjust C_v_ for accurate abundance estimation

A special case may occur in the cluster search process. When the sequencing data contains multiple strains in different clusters, some overlapping *k*-mers introduced during the CST optimization will cause FPs for the nodes’ *k*-mer matches. Specifically, suppose we reach a reconstructed node v after a leaf *r* has been identified to be present with an estimated abundance *A_r_*. In this case, the match counts of the *k*-mers in *K_v_* ⋂ *L_r_* are the sum of *k*-mer match counts from both *r* and strains in *v^T^*. Using the estimated abundance *A_r_*, we can estimate the match counts of *K_v_* ⋂ *L_r_* originated from *r* based on the Poisson distribution with λ = *A_r_*:

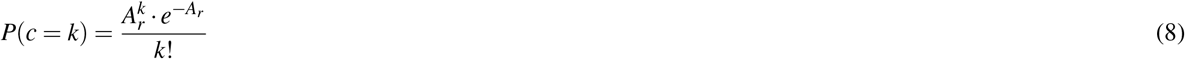

The estimated *k*-mer match vector from *r* is denoted as 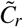 (the size is |K_v_ ⋂ L_r_|). Then, both of the vectors *C_v_* and 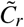 are sorted on the basis of increasing value of **k**-mer match count *c_i_*, and we adjust *C_v_* by 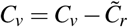. This method can be applied to multiple identified leaf nodes at different depths. After adjusting *C_v_* for all identified leaf nodes, we can use *C_v_* to calculate scoring metrics without any ambiguity.

### Choose *k*-mers for strain identification within the clusters

The CST is optimized for distinguishing clusters with similarity below a given cutoff. Using CST to distinguish highly similar strains can lead to a large number of weak nodes that cannot be augmented because of the large percentage of shared *k*-mers. Thus, once we pinpoint a cluster, we need a fine-grained method to distinguish highly similar strains. Once we pinpoint a cluster, the number of strains to distinguish is significantly reduced compared to the original problem space. Thus, we can afford to use all *k*-mers with distinguishing power. The first feature used is the unique *k*-mer from strain-specific regions, here we call it the strain-specific *k*-mer. The second feature used is the group-specific *k*-mer, which may come from structural variants (SVs) common to some strains. In a recent study^43^, SVs have been used to distinguish different strains. Inspired by that study, we extract group-specific *k*-mers from the SVs shared by some strains. However, relying only on strain-specific and group-specific *k*-mers still suffers from low resolution in some cases. For example, in Figure 12, both Strain4 and Strain5 have the same group-specific *k*-mers, and when the strain-specific *k*-mer of Strain5 is not present in the sample, we cannot make a fine distinction between the two strains. Therefore, to further improve the resolution, we need new features. SNVs and indels (insertions and deletions) between all strains are already applied to distinguish different strains in many studies^20, 29, 44^. Using these SNVs as well as indels, we can obtain unique *k*-mer sets that can distinguish all strains, and here we call these *k*-mers as *joint *k*-mers*. Thus, the third feature used is the joint *k*-mer, which contains SNVs and indels from core genomic regions present in all genomes. As shown in Figure 12, for all joint *k*-mers, although each *k*-mer is not strain-specific, the joint *k*-mer set for each strain is unique. However, the number of joint *k*-mers is often not as many as the first two types of *k*-mer (Supplementary Table S2), so they need to be combined together to improve the resolution of identification. By utilizing these three types of *k*-mers, we improve the resolution of identification and reduce the search space at the same time.

**Figure 12.**
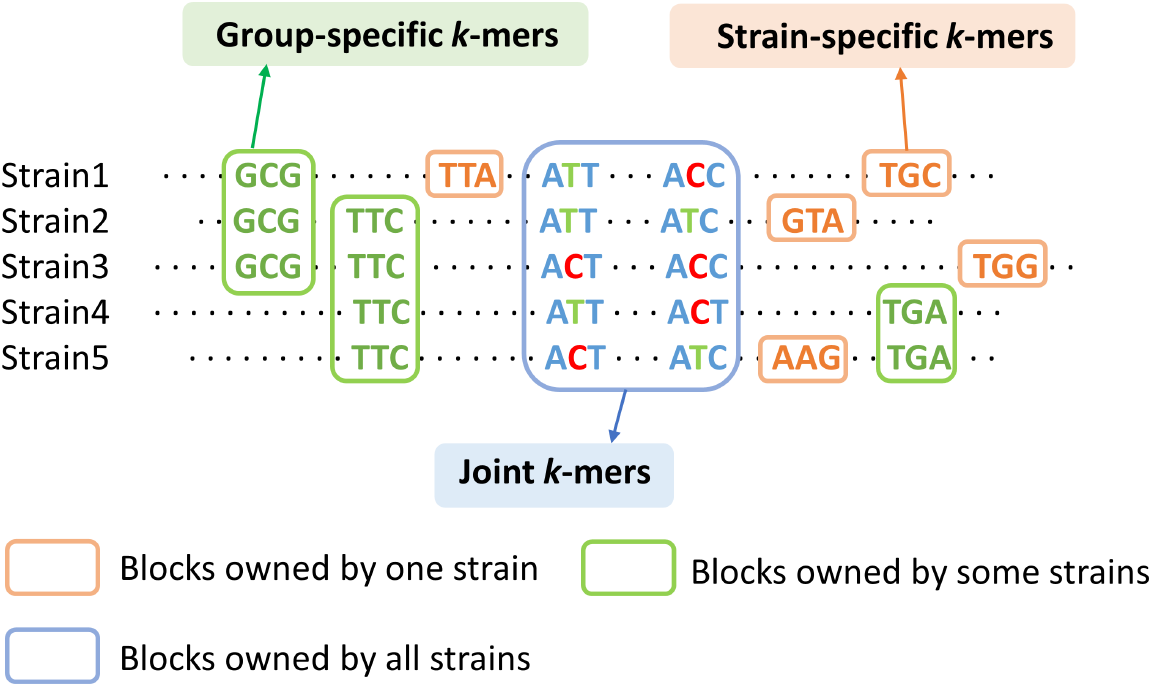
Use strain-specific *k*-mers, group-specific *k*-mers and joint *k*-mers to distinguish five strains in the same cluster. Each strain has a unique *k*-mer combination.

To effciently extract these *k*-mers, we utilize Sibeliaz^45^, an efficient tool designed for identifying locally collinear blocks in closely related genomes. Based on the blocks generated by Sibeliaz, we have developed an hash-based algorithm to extract these *k*-mers from strain genomes efficiently and save them in a matrix for later usage. The algorithm’s main pseudocode is shown in Supplementary Section 1.1. The input of the algorithm is strain genomes within the same cluster and blocks generated by Sibeliaz. By using the efficient hash table, the algorithm can extract target *k*-mers quickly. Finally, all extracted *k*-mers will be saved in a sparse matrix. Denote this matrix as a *M × N* sparse matrix. This matrix’s *M* rows represent *k*-mers found by the algorithm and *N* columns represent strains in the same cluster. In this matrix, a cell *X_k,i_* is 1 if the strain *i* possesses *k*-mer *k*. Otherwise, it is 0.

### Identify strains using chosen *k*-mers

After extracting the *k*-mers in the previous step, we need to use these features for strain identification. There are two main issues. One question is which strains are present and the second question is what is the relative abundance of these strains. To disentangle complex communities of closely related strains in the same cluster, we apply the iterative matrix multiplication to determine all coexisting strains and predict their relative abundance using elastic net regression.

The main goal of the iterative matrix multiplication is to determine strains in the same cluster by using three types of *k*-mers (Figure 12) in the *k*-mer matrix. To achieve this goal, we compare the *k*-mers in the sample to those of the *k*-mer matrix using an iterative strategy similar to that of QuantTB^28^. The method is described as follows. Given the cluster selected by the tree search and *k*-mers from its *k*-mer matrix X, we will apply Jellyfish^42^ to count all these selected *k*-mers in the sequencing data. Denote the occurrences for all selected *k*-mers from the Jellyfish as *M*-vector *y*: *y* = (*y*_1_,*y*_2_,*y*_3_,…,*y_M_*)^T^, where *y_i_* ≥ 0 and represents the occurrences of ith *k*-mer in the matrix. However, the overlapping *k*-mers from other identified clusters could lead to false *k*-mer matches or wrong abundance estimation. To remove the influence of other clusters, if one *k*-mer is found in other clusters detected by the tree search, then its occurrence will be replaced with 0. For the *M* × *N k*-mer matrix *X*, it can be written as *X* = (*s*_1_, *s*_2_, *s*_3_,…,*s_N_*), and the *k*-mer vector *s_j_* is defined as:

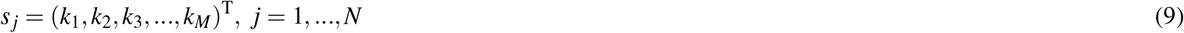

where 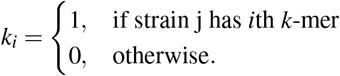, and N is the number of strains in the matrix.

Based on *X* and *y*, we use the iterative matrix multiplication, which can detect all possible strains in a sample accurately and quickly. Given *X* and y, the function will calculate a score *f_j_* = *s_j_* · *y* for each strain. It’s noted we regard values beyond the range of the 5nd and 95th percentile to be outliers, and we will set the value of all outliers to 0. The function will rank all strains according to their scores. After ranking, the function will output the top 1 strain in the ranked list and then update *y* by replacing the occurrences of all *k*-mers in identified strain with 0. Then the function will repeat this process. It continues to calculate the score and identify the best-matched strain in each iteration until the occurrences of *k*-mers with nonzero value is lower than the given threshold. The default threshoold is 31 * 40 = 1240, which means a sufficient number of SNVs (40) still remain in the sample and in the database for reliable classification, and this parameter can be changed by the user.

Knowing the possible strains in the sample, we use the elastic net regression model to predict sequencing depths and relative abundances of identified strains. Here, we choose the elastic net model instead of the lasso model because the former could better fit the abundance profile among strains compared to the method based on the lasso model like Strainest (Figure 4), and the lasso model tends to underestimate the number of strains, leading to a decrease in recall (Figure 3A). After iterative matrix multiplication, we obtain the filtered *k*-mer matrix *X*′ = *M* × *N*′, where *N*′ is the number of identified strains. Sequencing depths, which are the regression coefficients *β*′, are predicted minimizing the elastic net penalized residual sum of squares:

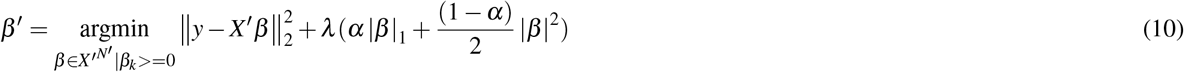

where the regression coefficients *β*′ should be nonnegative values. α and λ are two important parameters that will affect the model performance and therefore need to be tuned. We have designed a function to tune the α and λ based on crossvalidation to obtain the model with the lowest predictive error. Given this best model, we calculate the strain relative abundance *a* = (*a*_1_, *a*_2_, *a*_3_,…, *a_N_*′) by normalizing the regression coefficients *β*′ of the model. However, if multiple clusters are detected by the tree search, the relative abundance of one strain *i* will be recalculated according to the abundance of clusters. So, the final relative abundance (RA) of each strain *i* is calculated as:

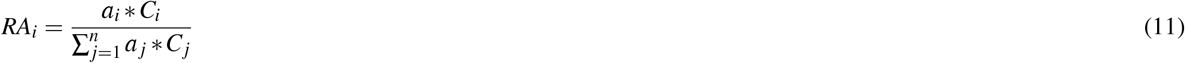

where *C* is the abundance of the cluster predicted by the tree search, and *n* is the total number of all identified strains.

### Prediction accuracy evaluation

In order to test the performance of each method, we calculated the recall, precision, and F1 score for every test category. True positive (TP) refers to the number of correctly identified strains. False negative (FN) refers to the number of strains present in the sample that were not identified. The definition of false positive (FP) can be found in the first paragraph of Results section. Some examples for illustrating these metrics are given in Supplementary Figure S2.

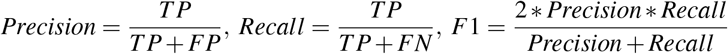

In all experiments, we used the Jensen-Shannon divergence (JSD)^46^ to measure the distance between the true and predicted relative abundance (RA). If the predicted and true RA have different dimensions, we will calculate JSD by adding zeros to the one with the lower dimension. Suppose there are two probability distributions T and P, then their Jensen–Shannon divergence is a value between [0,1] and defined as:

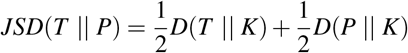

where

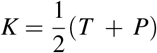

and *D*(*T* || *P*) is called the Kullback–Leibler divergence and it is defined as:

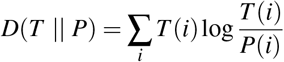

## FUNDING

This work is supprted by Hong Kong Research Grants Council (RGC) General Research Fund (GRF) 11206819 and 726 Hong Kong Innovation and Technology Fund (ITF) MRP/071/20X.

## Notes

### Competing Interest Statement

The authors have declared no competing interest.

