## Supplementary Data for "Accurate strain-level microbiome composition analysis from short reads"

### Supplementary information for "Accurate strain-level microbiome composition analysis from short reads"

Jan 2022

### 1 Supplementary Methods

#### 1.1 *k*-mer matrix construction

The pseudocode of constructing the k-mer matrix is given in Algorithm 1. To speed up strain search in the same cluster, we use a two-dimensional hash table $H$, which keeps track of the matched *k*-mers of each strain. $H$ has size of $m$ by $n$, where $m$ is the number of *k*-mers in the cluster and $n$ is the number of strains in the cluster. Each cell $H[i][j]$ (0 or 1) represents whether the ith *k*-mer is present in the jth strain. By using $H$, the algorithm can easily find strain-specific *k*-mers. Then, the algorithm divides the blocks obtained by Sibeliaz [1] into two parts. The first part is the blocks owned by some strains, which are SVs. The second part is the blocks that are shared by all strains, which are used to find joint *k*-mers. For *k*-mers from the second part, we use $H$ to keep all eligible *k*-mers as a way to obtain all joint *k*-mers. An eligible *k*-mer here means that the strains with this *k*-mer are a subset of all strains in the block.

**Algorithm 1** Build the *k*-mer matrix for strains in the same cluster.

**Require:** The strain genome set R $= \{R_1, R_2, \ldots, R_N\}$ with N strains. Collinear blocks $B$ of $R$ generated by Sibeliaz.
Define an empty sparse matrix $X$. ▷ The output matrix
Define a two-dimensional hash table $H$ for recording the strain label of all *k*-mers
Define a hash table $S$ for recording all output *k*-mers
**for** $R_1$ to $R_N$ **do** ▷ Initalize $H$
  **for** all *k*-mers $x \in R_i$ ($i = 1$ to $N$) **do**
    $H[x][R_i] = Null$, $H[revcomp(x)][R_i] = Null$ ▷ $revcomp$ returns the reverse compelement of a dna sequence.
**for** all *k*-mers $x \in H$ **do** ▷ Find strain-specific *k*-mers of each strain
  **if** $len(H[x]) == 1$ **then**
    $S[x] = Null$, $S[revcomp(x)] = Null$
**for** all blocks $b \in B$ **do**
  **for** all *k*-mers $x \in b$ **do**
    **if** not $len(b) == N$ **then** ▷ Find group-specific *k*-mers from blocks
      **if** $len(H[x]) == len(b)$ **then**
        $S[x] = Null$, $S[revcomp(x)] = Null$
    **else** ▷ Find joint *k*-mers from blocks
      **if** $H(x) \subset R$ **then**
        $S[x] = Null$, $S[revcomp(x)] = Null$
Initialize $X$ as a $N \times M$ matrix, $M = len(S)$
**for** all *k*-mers $x \in S$ **do** ▷ Fill out the *k*-mer matrix
  **for** $R_1$ to $R_N$ **do**
    **if** $R_i$ ($i = 1$ to $N$) $\in H[x]$ **then**
      $X[x, R_i] = 1$
    **else**
      $X[x, R_i] = 0$

### 2 Supplementary Figures

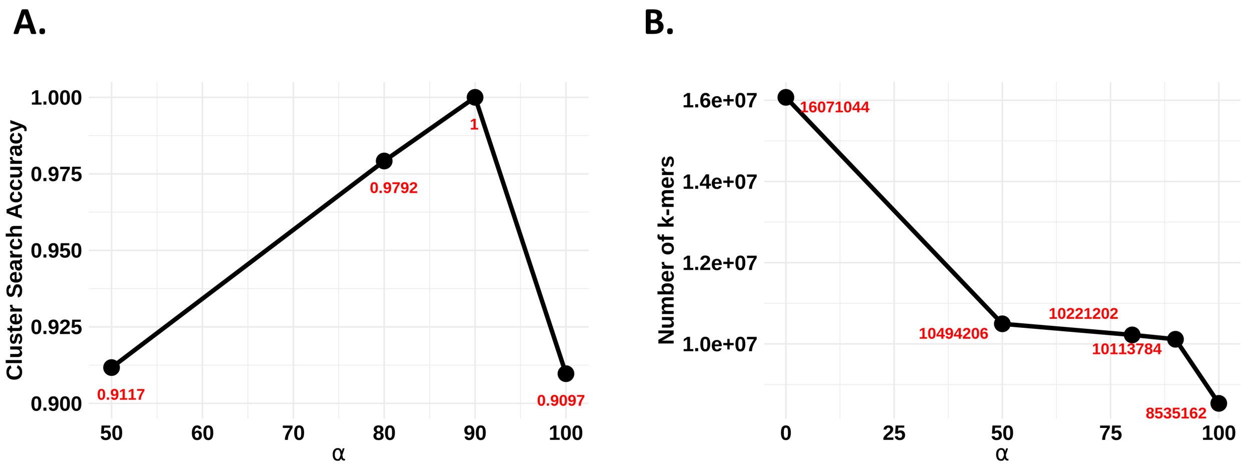


**Supplementary Figure S1.** (A). The cluster search accuracy when $\alpha$ is 50, 80, and 100, respectively. We generate 1,008 "single-strain" simulated datasets of S. epidermidis to test the cluster search performance. The experiments show that the performance is the best when $\alpha = 90$. When we only use $k$-mers that must occur in all strains ($\alpha = 100$), the resolution of the cluster search decreases. (B). The numbers of selected $k$-mers using different $\alpha$ values from a relatively large cluster that contains 201 *E.coli* strains.

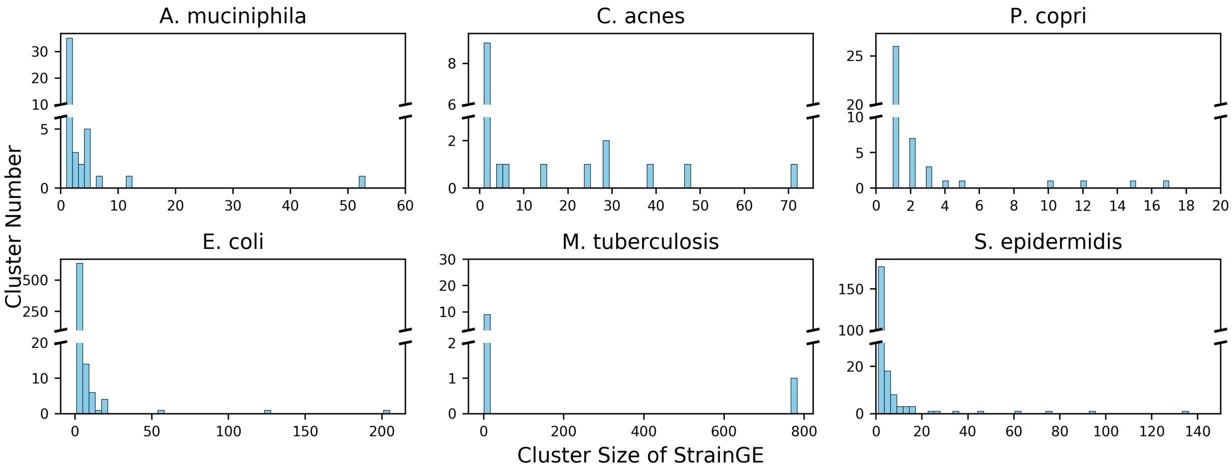


**Supplementary Figure S2.** The histogram of cluster sizes for clusters in the StrainGE's reference databases. For each cluster, only one representative strain will be selected and put into the final database for the identification. Although many clusters are small, there are also big ones. Using only the representative strain for the big clusters significantly decreased the resolution of strain identification.

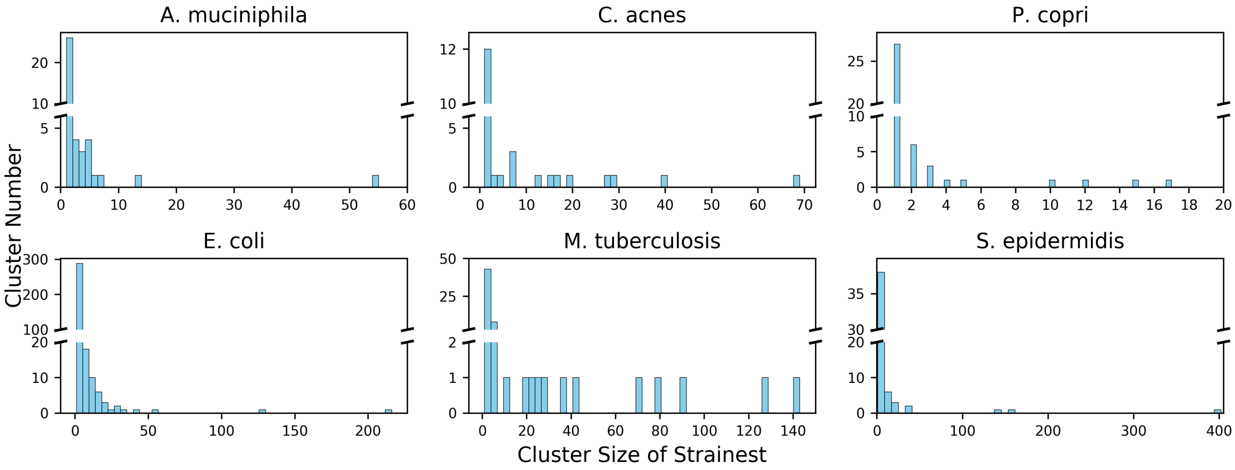


**Supplementary Figure S3.** The histogram of cluster sizes for clusters in Strainest's reference databases. For each cluster, only one representative strain will be selected and put into the final database for the identification. Although many clusters are small, there are also big ones. Using only the representative strain for the big clusters significantly decreased the resolution of strain identification.

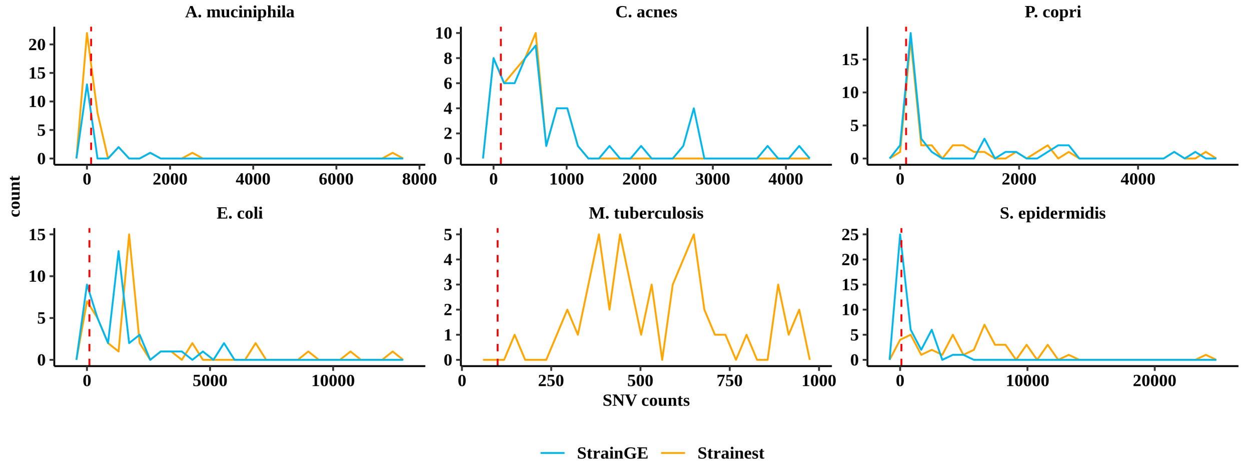


**Supplementary Figure S4.** The histogram of different SNVs between the representative strains and the actual strains identified by StrainGE and Strainest in the simulated "single-strain" datasets. The SNVs are detected by MUMmer [2]. The red line in the figure refers to 100 SNVs between the representative strain and the acutual strain.

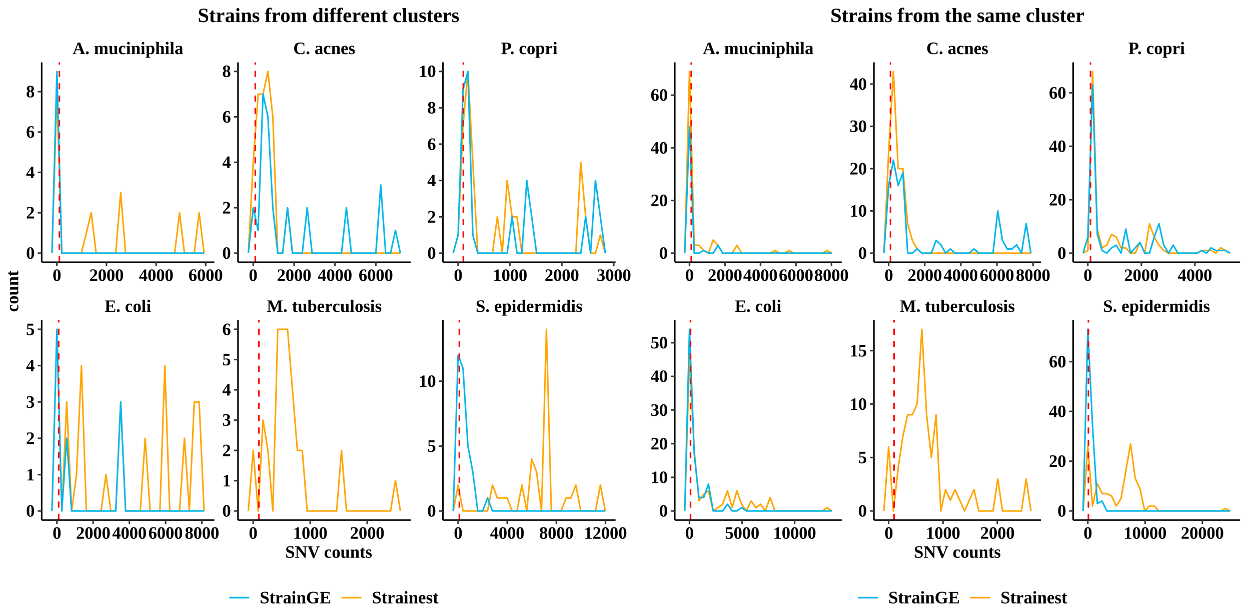


**Supplementary Figure S5.** The histogram of different SNVs between the representative strains and the actual strains identified by StrainGE and Strainest in the simulated "multiple-strain" datasets. The SNVs are detected by MUMmer [2]. The red line in the figure refers to 100 SNVs between the representative strain and the acutual strain.

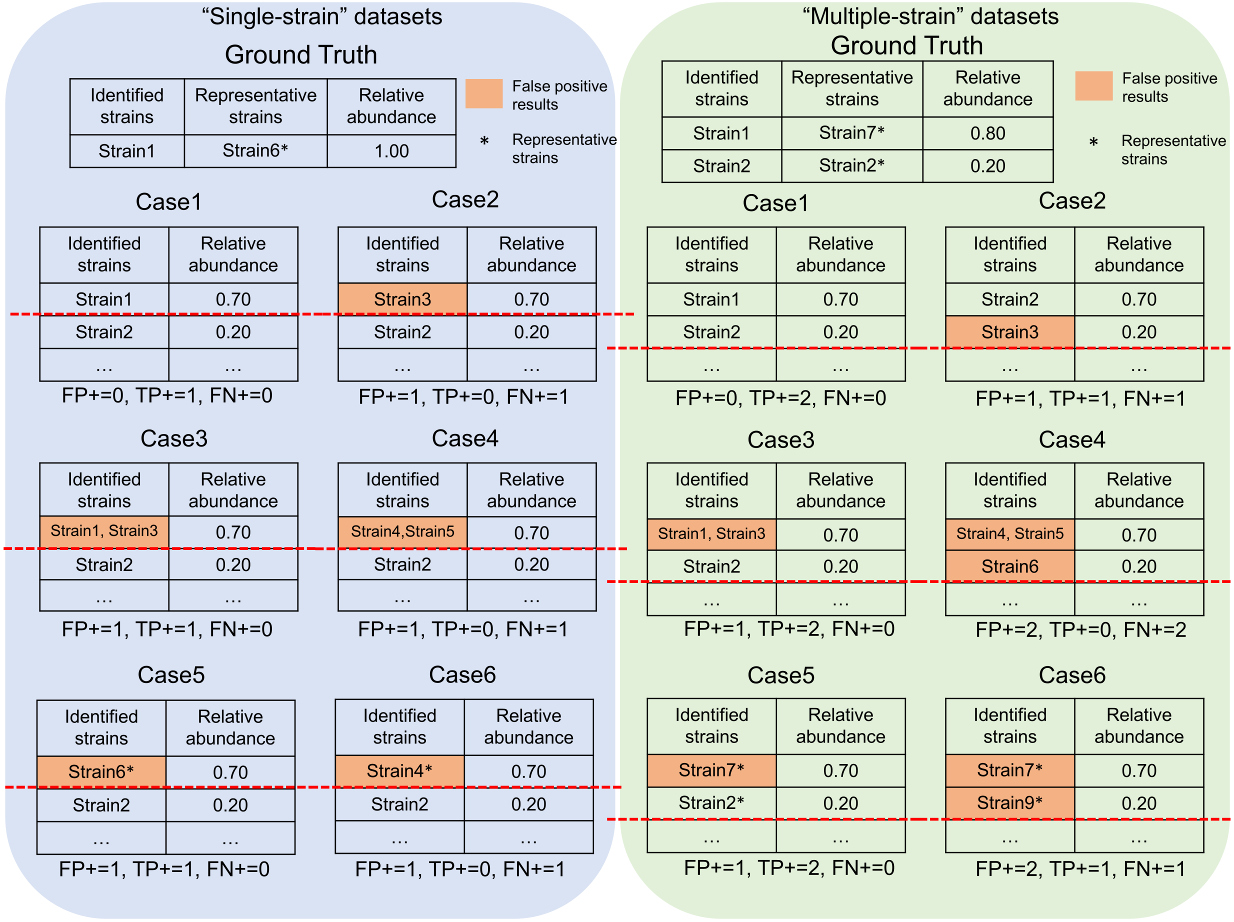


**Supplementary Figure S6.** TP, FP and FN in several cases. The red dash line in the figure indicates we only consider the top $x$ ranking strains when we calculate these metrics for the simulated datasets, where $x$ is the number of strains present in the simulated datasets.

#### 3 Supplementary Tables

| Species | # of strains | Before Clustering | | | After Clustering | | |
|---|---|---|---|---|---|---|---|
| | | # of $k$-mers | # of unique $k$-mers | # of strains without unique $k$-mers | # of $k$-mers* | # of increased unique $k$-mers | # of strains without unique $k$-mers |
| *E. coli* | 1,433 | 192,325,016 | 69,097,590 | 60 | 16,071,080 | 30,587,430 | 49 |
| *S. epidermidis* | 995 | 65,345,200 | 28,559,277 | 110 | 7,354,669 | 29,709,206 | 26 |
| *M. tuberculosis* | 792 | 42,221,860 | 29,004,912 | 0 | 30,011,796 | 9,985,111 | 0 |
| *C. acnes* | 275 | 16,923,331 | 4,107,627 | 74 | 6,245,590 | 1,845,459 | 72 |
| *A. muciniphila* | 157 | 45,911,791 | 10,862,010 | 17 | 6,488,987 | 1,501,606 | 12 |
| *P. copri* | 112 | 97,438,063 | 42,697,948 | 0 | 7,942,630 | 1,509,892 | 0 |

**Supplementary Table S1.** Comparison of the number of $k$-mers before and after clustering. "*" in the table means these $k$-mers are from the largest cluster after clustering. "# of increased unique $k$-mers" is obtained by summing the difference between the number of unique $k$-mers before and after clustering of each strain, where strains without unique $k$-mers before clustering are not included in the calculation.

| Species | # of strains | # of $k$-mers | # of chosen $k$-mers | # of strain-specific $k$-mers | # of group-specific $k$-mers | # of joint $k$-mers |
|---|---|---|---|---|---|---|
| *E. coli* | 201 | 16,071,080 | 6,614,270 | 1,990,030 | 3,365,764 | 1,258,476 |
| *S. epidermidis* | 92 | 7,354,669 | 3,024,846 | 1,396,638 | 1,232,400 | 395,808 |
| *M. tuberculosis* | 768 | 30,011,796 | 8,832,212 | 764,110 | 3,778,226 | 4,289,876 |
| *C. acnes* | 70 | 6,245,590 | 2,049,674 | 722,356 | 272,526 | 1,054,792 |
| *A. muciniphila* | 46 | 6,488,987 | 832,674 | 556,508 | 206,952 | 69,214 |
| *P. copri* | 17 | 7,942,630 | 108,408 | 26,670 | 37,722 | 44,012 |

**Supplementary Table S2.** The statistics of different types of $k$-mers in the largest cluster of each species.
